# Molecular determinants underlying functional divergence of TBP homologs

**DOI:** 10.1101/2025.10.03.680374

**Authors:** Jieying H. Cui 崔洁颖, Henry Young, Stephane Flibotte, Donglei Cai 蔡冬雷, Sheila S. Teves

## Abstract

The TATA-box binding protein (TBP) is a highly conserved basal transcription factor and a core component of the pre-initiation complex (PIC) for all three eukaryotic RNA polymerases (RNA Pols). Despite this conservation, TBP function diverges across species. In yeast, TBP is required for all three RNA Pols, whereas in mammals, it is essential only for RNA Pol III, but not RNA Pol I or II. To examine the evolutionary divergence of TBP homologs in different species, we determined the ability of murine TBP and its paralogs to complement endogenous TBP in *Saccharomyces cerevisiae*. Despite their highly conserved DNA-binding domains, murine TBP homologs were unable to fully rescue the lethality caused by TBP inactivation in yeast. This incomplete complementation correlates with a failure to fully support binding by RNA Pols II and III. Furthermore, we show that the divergent N-terminal domain (NTD) of TBP contributes to species-specific activity and modulates RNA Pol II binding changes in stress-induced response. Lastly, we demonstrate a negative correlation between the length of the intrinsically disordered NTD of TBP family proteins and the gene density in different species, suggesting that the NTD may have contributed to increasingly complex gene regulation during evolution.

## Introduction

As the first step in gene regulation, transcription initiation requires a stepwise assembly of the RNA Polymerase (RNA Pol) pre-initiation complex (PIC) at gene promoters (Matsui *et al*, 1980; Segall *et al*, 1980). Eukaryotes share three RNA Pols (Pol I-III), largely responsible for ribosomal, messenger, and transfer RNA (rRNA, mRNA, tRNA), respectively (Werner & Grohmann, 2011; Roeder & Rutter, 1969; Petes, 1979; Hurowitz & Brown, 2003; Dieci *et al*, 2013). Despite differences among the three PICs, the TATA-box binding protein (TBP) is the only general transcription factor required by all three (Kwon & Green, 1994; Cormack & Struhl, 1992b; White *et al*, 1992; Schröder *et al*, 2003; Shen *et al*, 1998). TBP contains a conserved DNA-binding core domain that recognizes the promoters and binds to the DNA minor groove, and a divergent N-terminal domain (NTD) (Starr & Hawley, 1991; Kim *et al*, 1993; Lee *et al*, 1991). TBP function is essential for viability and highly conserved across eukaryotes (Rowlands *et al*, 1994).

Since its discovery in 1988, TBP has been central to studies of eukaryotic transcription (Nakajima *et al*, 1988; Buratowski *et al*, 1988; Hoey *et al*, 1990). Although TBP essentiality initially hindered functional studies, conditional systems have revealed species-specific differences. In *Saccharomyces cerevisiae*, acute depletion of TBP via the anchor-away system causes genome-wide loss of RNA Pol II transcription (Petrenko *et al*, 2017, 2019), while temperature-sensitive (ts) mutants also dramatically impair RNA Pol I and Pol III transcription (Steffan *et al*, 1996; Cormack & Struhl, 1992b). By contrast, acute depletion of mouse and human TBP via the Auxin-inducible degron system does not affect RNA Pol I and Pol II occupancy in mammalian cells, but severely reduces RNA Pol III recruitment (Santana *et al*, 2022; Kwan *et al*, 2023, 2024; Cui *et al*, 2025). Divergence is also evident in the NTD of TBP. The NTD is dispensable for yeast cell growth (Cormack *et al*, 1994; Lee & Struhl, 2001) but essential for mouse embryonic viability and placental development (Hobbs *et al*, 2002).

Gene duplication has further diversified TBP function through the emergence of paralogs (Akhtar & Veenstra, 2011; Ravarani *et al*, 2020). For instance, the insect-specific TBP paralog called TBP-related factor 1 (TRF1) aids RNA Pol III transcription (Takada *et al*, 2000). Bilaterians acquired TRF2, which, in mice, is widely expressed and is crucial for spermiogenesis (Martianov *et al*, 2001; Zhang *et al*, 2001). Vertebrates have gained TRF3, which in mice mediates oocyte-specific transcription (Akhtar & Veenstra, 2009; Yu *et al*, 2020). Unlike yeast, animals have evolved multiple non-redundant TBP paralogs that may facilitate more diverse gene regulation functions.

We have previously shown that sequence similarity in the conserved core domain (40%-85%) does not predict function in mouse embryonic stem cells (mESCs) (Cui *et al*, 2025). Specifically, TRF2 and yeast TBP can bind to RNA Pol II and III promoters in mESCs, albeit at altered levels, whereas TRF3 cannot, despite its high similarity to mouse TBP (Cui *et al*, 2025). These molecular behaviors are reflective of the intrinsic differences in DNA binding dynamics, contributing to their species-or tissue-specific roles (Cui *et al*, 2025). Interestingly, the paralogs have diverged functionally from the mouse TBP even more than the yeast ortholog, raising the question of how mammalian homologs behave in yeast.

Here, we assessed how TBP homologs and their domains contribute to functional divergences. Using a conditional TBP depletion system in yeast, we tested whether different mouse TBP, the paralogs TRF2 and TRF3, and their truncations can functionally replace the endogenous TBP protein. Despite conservation of the core domain, mouse TBP homologs failed to fully rescue yeast TBP loss, correlating with reduced Pol II and III transcription. We further show that the NTD contributes to species-specific activity and stress-induced transcriptional responses. Finally, comparative analyses across >300 species revealed that NTD length scales with total TBP length and inversely with gene density, suggesting an evolutionary role in complex gene regulation.

## Result

### Differential rescue of TBP lethality in yeast cells by homologous TBPs

To generate a TBP replacement system, we used a previously established *S. cerevisiae* strain with a temperature-sensitive TBP allele (tsTBP), which is conditionally inactivated at the restrictive temperature of 36.5 ± 0.5 °C (Cormack & Struhl, 1992a). We introduced HA-tagged constructs expressing wild-type yeast TBP (HA-yTBP), mouse TBP (HA-mTBP), the paralogs TRF2 and TRF3 (TRF2-HA, TRF3-HA), truncations of the conserved core domains (HA-mTBPc, TRF3c-HA), and an empty vector control (Fig. S1A). All constructs were driven by the 5’ and 3’ UTR of the endogenous yeast TBP, and protein expression was verified by anti-HA Western blotting (Fig. S1B).

Cell growth and viability were assessed under permissive and restrictive temperatures using both spot assays (cumulative growth) and liquid culture growth curves (time-resolved changes) for 24 to 36 hours. At the permissive temperature (29 ± 0.5 °C), HA-mTBP, TRF2-HA, and HA-mTBPc grew comparably to HA-yTBP (Fig. 1A, S1C), reaching ∼85% of the wild-type control as measured by area under the curve (AUC) of the growth assays (Fig. 1B). In contrast, TRF3-HA and TRF3c-HA strains showed severe growth defects at the permissive temperature, even worse than the tsTBP empty vector strain, suggesting a dominant negative effect (Fig. 1A-B, S1C).

**Figure 1.**
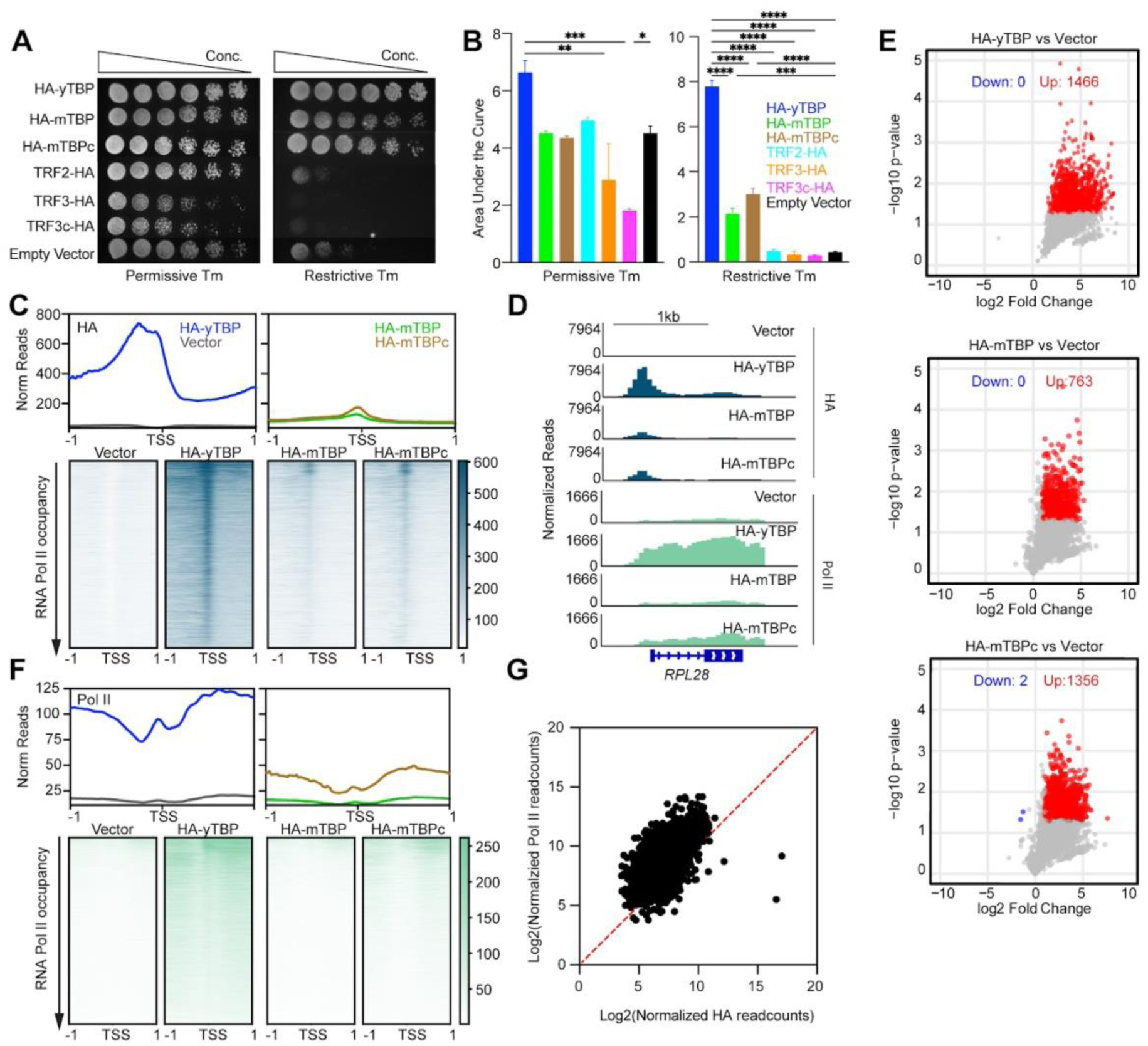
Differential rescue of TBP lethality in yeast cells by homologous TBPs. (A) Spot assay of parental ts-TBP strain expressing HA-yTBP, HA-mTBP, HA-mTBPc, TRF2-HA, TRF3-HA, TRF3c-HA and vector at permissive (left) and restrictive temperature (right). (B) Area under the curve of growth assay curves of the same strains as (A) with error bars representing the standard error of the mean (SEM). (C) Genome-wide average plots (top) and heatmaps (bottom) arranged by decreasing RNA Pol II occupancy of HA ChIP-seq for HA-yTBP and vector (left), HA-mTBP and HA-mTBPc (right) in a 2 kb window surrounding the transcription start site (TSS) of all genes. (D) Gene browser tracks at *RPL28* for HA and RNA Pol II ChIP-seq of vector, HA-yTBP, HA-mTBP and HA-mTBPc. (E) Volcano plot of HA ChIP-seq normalized reads as a fold-change of HA-yTBP (top), HA-mTBP (middle), and HA-mTBPc relative to vector (bottom) against the -log10 of p-value. (F) Genome-wide average plots (top) and heatmaps (bottom) arranged by decreasing RNA Pol II occupancy of RNA Pol II ChIP-seq for HA-yTBP and vector (left), HA-mTBP and HA-mTBPc (right) in a 2 kb window surrounding the TSS of all genes. (G) Normalized read counts of HA ChIP-seq signal in the promoter (−50 to TSS) vs. RNA Pol II ChIP-seq in the gene body of each Pol II gene. Statistics calculated using one-way ANOVA: *** p-value ≤ 0.001, **** p-value ≤ 0.0001. Non-significant statistics is not shown.

At the restrictive temperature, the empty vector strain was non-viable, confirming successful inactivation of the tsTBP (Fig. 1A-B, S1C). HA-mTBP and HA-mTBPc provided partial rescue at the restrictive temperature, achieving ∼25% of HA-yTBP growth, whereas TRF2-HA and TRF3-HA failed to rescue (Fig. 1A-B, S1C). Despite 90% similarity between TRF3 and mTBP core domains, TRF3c-HA also failed to restore viability (Fig. 1A-B, S1C).

To examine DNA binding, we performed spike-in normalized HA ChIP-seq in two biological replicates after 3 hours at the restrictive temperature, with expression confirmed by Western blotting (Fig. 1C-D, S1B, S1D-E). After alignment and de-duplication, spike-in normalized reads from averaged and individual replicates were displayed as average profiles and heatmaps in a 2kb window around the transcription start site (TSS) for all genes (Fig. 1C, S1E). HA-yTBP bound strongly at promoters of expressed genes, whereas the empty vector showed background signal (Fig. 1C). Both HA-mTBP and HA-mTBPc bound RNA Pol II promoters, though globally at only ∼28% HA-yTBP levels (Fig. 1C). Notably, HA-mTBPc displayed a modestly higher promoter occupancy than HA-mTBP, exemplified at the *RPL28* locus (Fig. 1C-D). Quantification relative to the empty vector identified 763 and 1356 significantly bound sites, respectively (Fig. 1E), suggesting a modest inhibitory effect by the mouse NTD. Gene Ontology (GO) analysis of the HA-mTBPc-versus HA-yTBP-bound sites identified the same pathways, including ribosome biogenesis and processing (Fig. S1F), suggesting that the binding of the mouse paralogs is governed by wild-type yeast TBP binding levels.

We next examined their ability to recruit RNA Pol II itself. Spike-in normalized Rpb3 ChIP-seq was performed on cells grown at the restrictive temperature for 3 hours, with averaged and individual replicates showing that tsTBP inactivation in the empty vector caused ∼90% loss of RNA Pol II occupancy relative to HA-yTBP (Fig. 1F, S1G). Although HA-mTBP bound to promoters weakly, it did not recruit RNA Pol II above background levels (Fig. 1D, F). In contrast, HA-mTBPc recruited ∼40% as much RNA Pol II as HA-yTBP (Fig. 1D, F). Furthermore, promoter occupancy by HA-mTBPc correlated with RNA Pol II binding (Fig. 1G), and genes with >4-fold higher RNA Pol II occupancy compared to empty vector were enriched for GO terms involved in the protein synthesis pathway, consistent with where HA-mTBPc is most bound (Fig. S1F, H). Taken together, these results suggest that the mouse core domain retains conserved functions of TBP in yeast, but the NTD reduces binding efficiency and Pol II recruitment, highlighting its role in species-specific activity.

### Molecular behavior of homologous TBPs in RNA Pol I and III genes

Next, we questioned how the orthologs bind to rRNA and tRNA genes mainly transcribed by RNA Pol I and III, respectively. A challenge is that the standard *S. cerevisiae* sacCer3 genome masks tandemly repeated rDNA sequences (Kim, 2006). Each transcription unit is separated by the Intergenic Spacer (IGS) and is composed of Promoter repeats, External and Internal Transcribed Spacers (ETS and ITS respectively), ribosomal RNA genes (18S, 5.8S and 25S) and the 5S rRNA gene (Kim, 2006). Therefore, we generated a customized genome in which rDNA repeats were collapsed into a single annotated rDNA “chromosome” (George *et al*, 2023), and aligned spike-in normalized HA ChIP-seq reads (Fig. 2A, S2A). We then quantified binding at the 35S promoter, the Pol I recruitment site. As expected, HA-yTBP showed robust binding, while the empty vector strain displayed background levels. HA-mTBP and HA-mTBPc retained binding at ∼45% and ∼60% of HA-yTBP levels, respectively (Fig. 2A-B), indicating that mouse TBP preserves partial activity at Pol I-transcribed genes.

**Figure 2.**
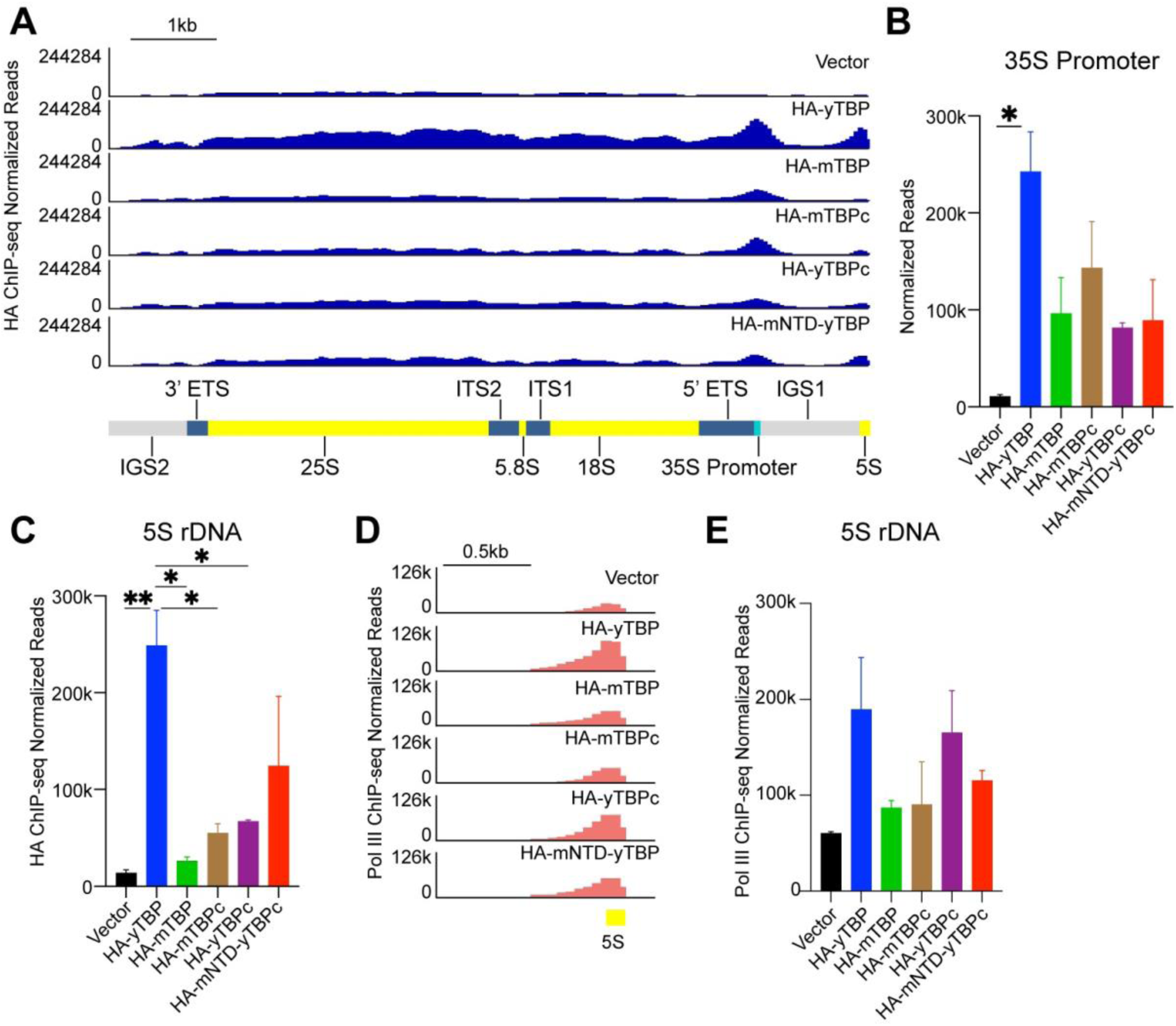
Molecular behavior of homologous TBPs in RNA Pol I genes. (A) Gene browser tracks of the custom rDNA chromosome (chrR) for HA ChIP-seq normalized reads of Vector, HA-yTBP, HA-mTBP, HA-mTBPc, HA-yTBPc and HA-mNTD-yTBPc. (B-C) Averaged normalized read counts of HA ChIP-seq at 35S promoter (B) and 5S rDNA (B). (D) Gene browser tracks of the custom chrR on 5S rDNA for RNA Pol III ChIP-seq normalized reads of Vector, HA-yTBP, HA-mTBP, HA-mTBPc, HA-yTBPc and HA-mNTD-yTBPc. (E) Averaged normalized read counts at 5S rDNA for RNA Pol III ChIP-seq data of all samples mentioned in (D). Error bars represent standard deviation of two biological replicates. Statistics calculated using one-way ANOVA: * p-value ≤ 0.1, ** p-value ≤ 0.01. Non-significant statistics is not shown.

The 5S rRNA gene, embedded in the rDNA array, is instead transcribed by Pol III using a Class I promoter (Paule, 2000; Schramm & Hernandez, 2002; French *et al*, 2008). Intriguingly, HA-mTBP and HA-mTBPc showed markedly reduced binding relative to HA-yTBP, with HA-mTBPc binding slightly more strongly than full-length HA-mTBP (Fig. 2A, C), consistent with an inhibitory role of the mouse NTD.

To test whether TBP homologs could recruit Pol III, we tagged the endogenous Pol III subunit RPC160 with Flag, validated expression by Western blot (Fig. S2B), and performed spike-in normalized ChIP-seq two biological replicates. Pol III occupancy at the 5S rDNA locus was robust with HA-yTBP but nearly absent in the empty vector strain. Neither HA-mTBP nor HA-mTBPc supported Pol III recruitment above background, despite detectable HA-mTBPc binding (Fig. 2D-E, S2C).

We next analyzed Pol III targets beyond rDNA, focusing on tRNA genes with Class II promoters (Fig. 3A-B, S3A) (Schramm & Hernandez, 2002). HA-yTBP bound strongly at tRNA promoters, exemplified by the *TRN1* tRNA-Pro gene, whereas the empty vector showed background levels (Fig. 3A). Genome-wide at all tRNA genes, HA-yTBP enrichment was ∼300-fold higher than the empty vector control (99.6% decrease to HA-yTBP) (Fig. 3B). In contrast, HA-mTBP and HA-mTBPc showed only ∼5-fold (98.8% decrease) and ∼8-fold (98% decrease) increases relative to the empty vector, respectively (Fig. 3A-B), with HA-mTBPc again binding modestly more strongly than full-length TBP (Fig. 3B-C). Notably, these signals were two orders of magnitude lower than those of HA-yTBP (Fig. 3B).

**Figure 3.**
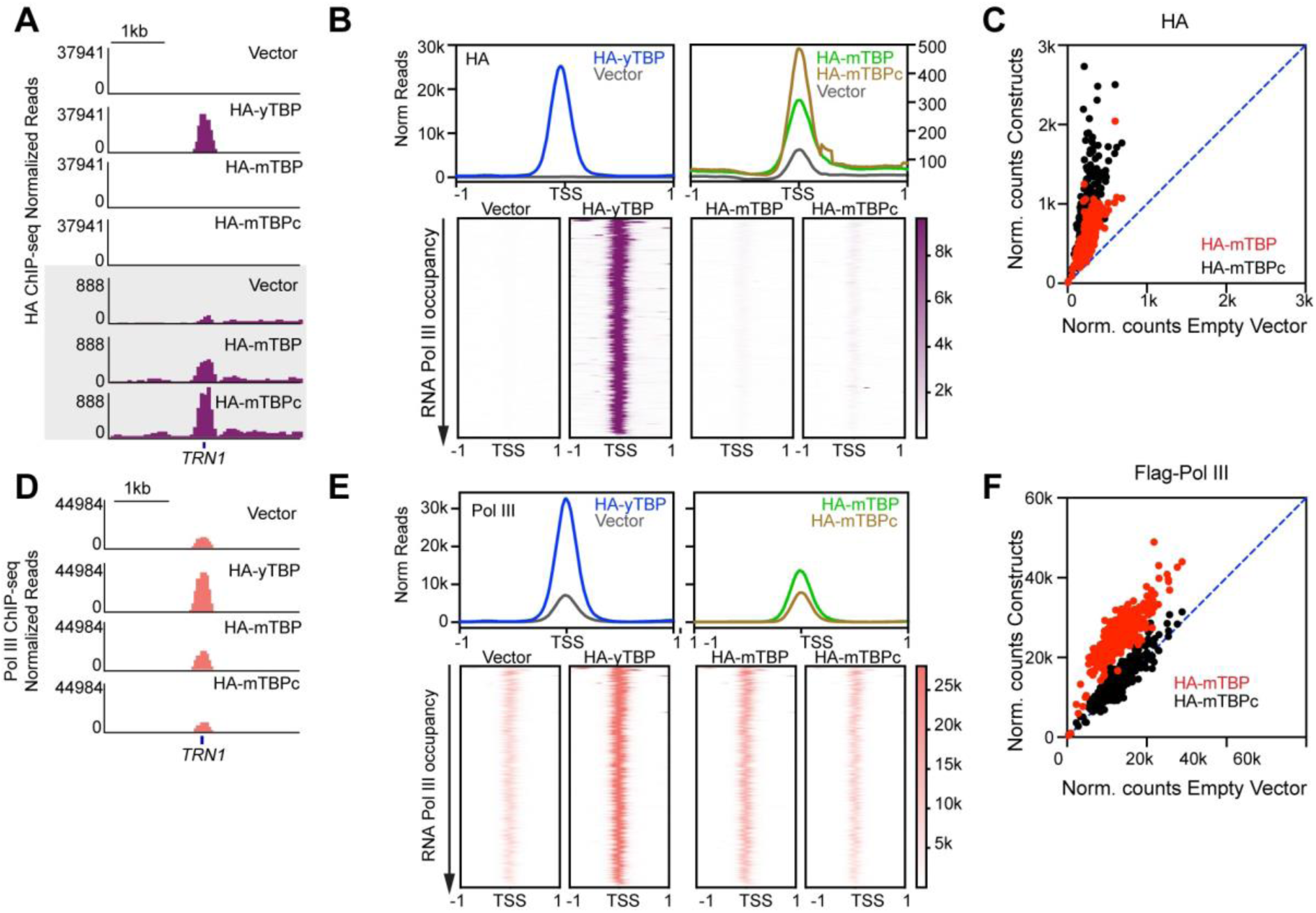
Molecular behavior of homologous TBPs in RNA Pol III genes. (A) Gene browser tracks at *TRN1* for HA ChIP-seq of vector, HA-yTBP, HA-mTBP and HA-mTBPc in two y-axis ranges. (B) Genome-wide average plots (top) and heatmaps (bottom) arranged by decreasing RNA Pol III occupancy of HA ChIP-seq for HA-yTBP and vector (left), HA-mTBP and HA-mTBPc (right) in a 2 kb window surrounding the TSS of all tRNA genes. (C) Normalized read counts of HA ChIP-seq signal of HA-mTBP and HA-mTBPc versus vector in the gene body (–50 to TES) region of each tRNA gene. (D) Gene browser tracks at *TRN1* for RNA Pol III ChIP-seq of vector, HA-yTBP, HA-mTBP and HA-mTBPc. (E) Genome-wide average plots (top) and heatmaps (bottom) arranged by decreasing RNA Pol III occupancy of RNA Pol III ChIP-seq for HA-yTBP and vector (left), HA-mTBP and HA-mTBPc (right) in a 2 kb window surrounding the TSS of all tRNA genes. (F) Normalized read counts of RNA Pol III ChIP-seq signal of HA-mTBP and HA-mTBPc vs. vector in the gene body (–50 to TES) region of each tRNA gene.

RNA Pol III normalized ChIP-seq reads at tRNA genes reflected the reduced TBP binding. Robust recruitment occurred in HA-yTBP strains but was strongly reduced in the empty vector and nearly absent with HA-mTBP or HA-mTBPc (Fig. 3D-F, Fig. S3B). Together, these results indicate that while mouse TBP homologs retain partial binding activity at Pol I-transcribed genes, they are severely impaired in binding and recruiting Pol III, underscoring the heightened sensitivity of Pol III to TBP divergence.

### The variable NTD differentially affects TBP binding and RNA Pols recruitment under homeostasis

To further investigate the divergent and disordered NTD of TBP, we generated an HA-tagged truncated yeast TBP core construct (HA-yTBPc) and a chimeric protein in which the mouse NTD was fused to the yeast TBP core (HA-mNTD-yTBPc). Both were introduced into the ts-TBP yeast strain and verified by Western blotting (Fig. S1D). Alongside the HA-yTBP and empty vector controls, we first assessed growth phenotypes by liquid culture and spot assays. Neither NTD deletion nor addition of the mouse NTD altered growth at permissive or restrictive temperatures (Fig. 4A-B, S4A), suggesting that the yeast core domain is the primary determinant of cell viability.

**Figure 4.**
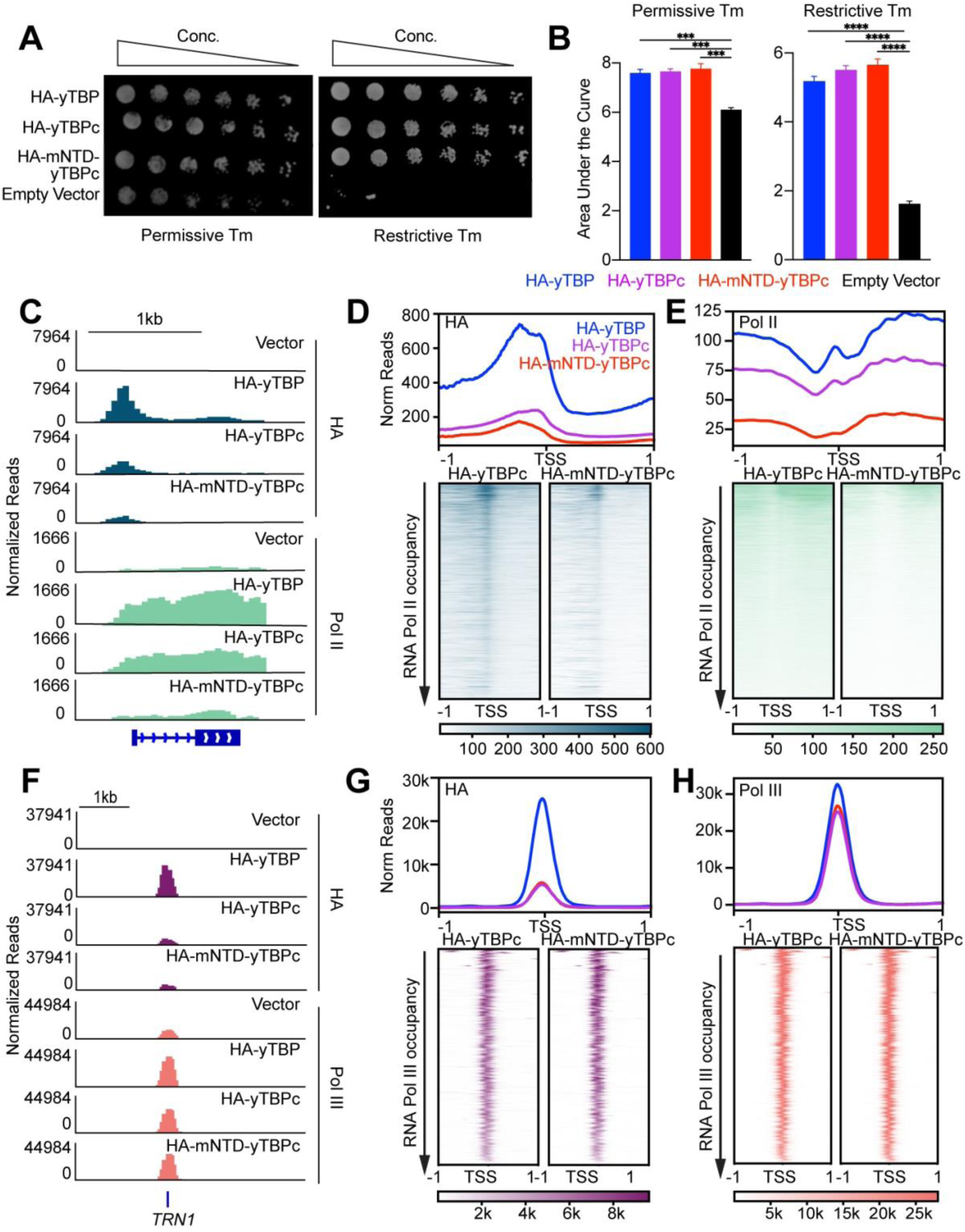
The variable NTD differentially affects TBP binding and RNA Pols recruitment under homeostasis. (A) Spot assay of parental ts-TBP mutant expressing HA-yTBP, HA-mTBP, HA-mTBPc, TRF2-HA, TRF3-HA and TRF3c-HA at the permissive (left) and restrictive temperature (right). (B) Area under the curve of growth assay curves of the same strains as (A) with error bars representing the SEM. (C) Gene browser tracks at *RPL28* for HA and RNA Pol II ChIP-seq of vector, HA-yTBP, HA-yTBPc and HA-mNTD-yTBPc. (D) Genome-wide average plots (top) and heatmaps (bottom) arranged by decreasing Pol II occupancy of HA ChIP-seq for HA-yTBP, HA-yTBPc and HA-mNTD-yTBPc in a 2 kb window surrounding the TSS of all genes. (E) Genome-wide average plots (top) and heatmaps (bottom) arranged by decreasing Pol II occupancy of RNA Pol II ChIP-seq for HA-yTBP, HA-yTBPc and HA-mNTD-yTBPc in a 2 kb window surrounding the TSS of all genes. (F) Gene browser tracks at *TRN1* for HA and RNA Pol III ChIP-seq of vector, HA-yTBP, HA-yTBPc and HA-mNTD-yTBPc. (G) Genome-wide average plots (top) and heatmaps (bottom) arranged by decreasing RNA Pol III occupancy of HA ChIP-seq for HA-yTBP, HA-yTBPc and HA-mNTD-yTBPc in a 2 kb window surrounding the TSS of all tRNA genes. (H) Genome-wide average plots (top) and heatmaps (bottom) arranged by decreasing RNA Pol III occupancy of RNA Pol III ChIP-seq for HA-yTBP, HA-yTBPc and HA-mNTD-yTBPc in a 2 kb window surrounding the TSS of all tRNA genes. Statistics calculated using one-way ANOVA: **** p-value ≤ 0.0001. Non-significant statistics is not shown.

We next examined the molecular consequences of NTD manipulation by performing anti-HA ChIP-seq in two biological replicates with spike-in normalization (Fig. 4C-D, S4B). At the *RPL28* locus, both HA-yTBPc and HA-mNTD-yTBPc displayed markedly reduced promoter occupancy compared to full-length HA-yTBP (Fig. 4C). Genome-wide, this decrease corresponded to ∼25% and ∼20% of HA-yTBP occupancy, respectively (Fig. 4D), revealing that the NTD contributes to global TBP binding capacity despite no overt effect on growth.

We then assessed RNA Pol II binding by performing two independent replicates of spike-in normalized Rbp3 ChIP-seq, with the results visualized for averaged and individual replicates (Fig. 4E, S4C). Pol II occupancy closely paralleled HA binding patterns both at RPL28 and genome-wide (Fig. 4C-E). Relative to HA-yTBP, the HA-yTBPc strain exhibited a ∼25% reduction in Pol II recruitment, while the HA-mNTD-yTBPc strain showed a ∼75% reduction, consistent with a stronger inhibitory influence of the mouse NTD on yeast TBP core activity (Fig. 4D-E). Notably, lack of the yeast-specific NTD also inhibited TBP and RNA Pol II binding, further supporting a species-specific role of the NTD in the regulation of TBP activity.

We next examined Pol I and Pol III behavior using the custom rDNA chromosome. Both HA-yTBPc and HA-mNTD-yTBPc displayed approximately half of the binding level of the full-length yeast TBP at the 35S rDNA promoter and 5S rDNA gene loci (Fig. 2A-C), consistent with the decreased binding capacity observed at Pol II promoters. For Pol III-transcribed tRNA genes, HA-yTBPc and HA-mNTD-yTBPc exhibited ∼80% reduced binding at TRN1 and globally (Fig. 4F-G, S4D). Surprisingly, RNA Pol III recruitment, measured by ChIP-seq of the Flag-tagged subunit, remained largely unaffected at *TRN1* and across the genome (Fig. 4F, H, S4E). This capacity to maintain Pol III occupancy despite diminished TBP binding likely accounts for the absence of a growth defect.

### The variable NTD modulates transcriptional responses to stress

Next, we investigated the functional role of the variable NTD under stress conditions in cell growth and transcriptional reprogramming. We performed growth assays for the ts-TBP strains expressing empty vector, HA-yTBP, HA-yTBPc, and HA-mNTD-yTBPc in media containing an oxidative stressor, diamide, at both permissive and restrictive temperatures (Fig. 5A, S5A). Diamide treatment in yeast triggers a transcriptional reprogramming that suppresses growth-related transcription and induces antioxidant and stress-defence gene expression (Gasch *et al*, 2000). At the permissive temperature, all strains exhibited similar growth phenotypes under diamide stress (Fig. 5A, S5A). However, at the restrictive temperature, both the truncated and chimeric constructs showed slower growth, quantified by area under the curve, relative to the full-length yeast TBP (Fig. 5A, S5A).

**Figure 5.**
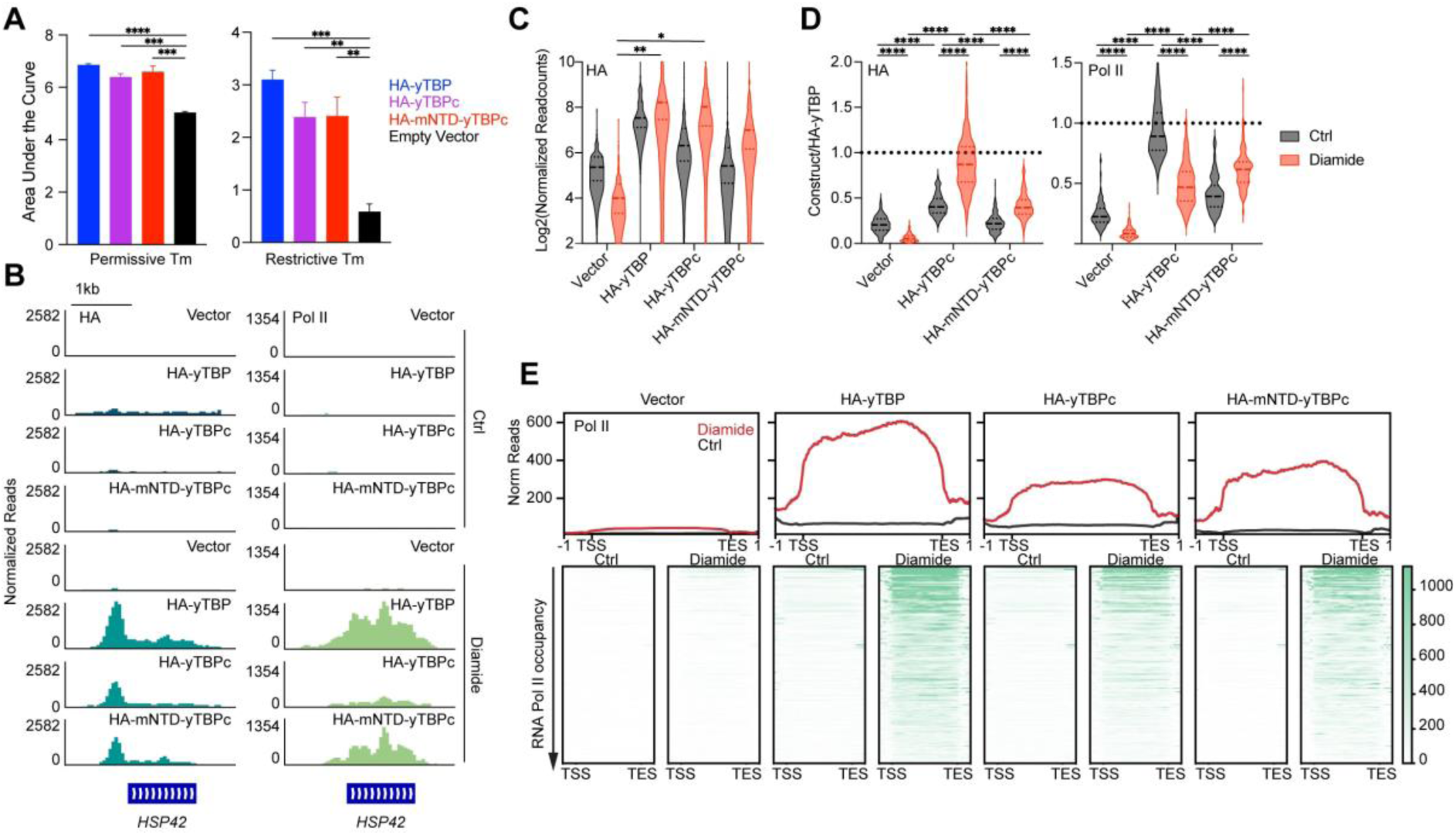
The variable NTD modulates transcriptional responses to stress. (A) Area under the curve of growth assays of the parental ts-TBP strain expressing HA-yTBP, HA-yTBPc, HA-mNTD-yTBPc and empty vector treated with 0.75 mM diamide at the permissive (left) and restrictive temperature (right), with error bars representing the SEM. (B) Gene browser tracks at *HSP42* for HA (left) and RNA Pol II (right) ChIP-seq of vector, HA-yTBP, HA-yTBPc and HA-mNTD-yTBPc at restrictive temperature only (Ctrl) and at restrictive temperature with diamide. (C) Log2 of normalized read counts of HA ChIP-seq signal in vector, HA-yTBP, HA-yTBPc and HA-mNTD-yTBPc in the promoter (−50 to TSS) of each diamide-induced RNA Pol II gene at both non-diamide (Ctrl) and diamide-treated conditions (Diamide), visualized as a violin plot. (D) The degree of induction was quantified by the ratio of vector, HA-yTBPc or HA-mNTD-yTBPc over HA-yTBP for all diamide-induced RNA Pol II genes at both non-diamide (Ctrl) and diamide-treated condition (Diamide), visualized as a violin plot. (E) Metagene plot (top) and heatmaps (bottom) arranged by decreasing RNA Pol II occupancy of RNA Pol II ChIP-seq for vector, HA-yTBP, HA-yTBPc and HA-mNTD-yTBPc from TSS to TES of diamide-induced genes at both non-diamide (Ctrl) and diamide-treated condition (Diamide). Statistics calculated using one-way ANOVA: *p-value ≤ 0.1, **p-value ≤ 0.01,*** p-value ≤ 0.001; **** p-value ≤ 0.0001. Non-significant statistics is not shown.

To probe molecular changes during stress, we performed spike-in normalized HA and RNA Pol II ChIP-seq after 3 hours of growth at the restrictive temperature and 15 minutes of diamide treatment, in two biological replicates (Fig. 5B, S5B). At the stress-induced gene *HSP42*, HA-yTBP binding increased ∼7-fold and RNA Pol II binding ∼18-fold upon diamide treatment compared to untreated controls, and more than 20-fold relative to empty vector controls (Fig. 5B). HA-yTBPc and HA-mNTD-yTBPc also showed increased binding at *HSP42*, but at ∼50% of HA-yTBP levels (Fig. 5B). Genome-wide, we identified 146 diamide-induced genes with ≥4-fold higher RNA Pol II binding in HA-yTBP strains compared to untreated controls. These genes were enriched for GO terms related to oxidative stress responses and protein refolding, consistent with known diamide responses (Fig. S5C).

Violin plots of HA ChIP-seq read counts at promoters (−50 to TSS) confirmed that both HA-yTBPc and HA-mNTD-yTBPc displayed increased binding at induced genes upon diamide treatment, but at lower levels than HA-yTBP (Fig. 5C). This reduced induction was reflected in the log2 ratio of HA occupancy, which was <1 for both truncated and chimeric proteins compared to full-length TBP (Fig. 5D, left). Interestingly, HA-yTBPc showed higher promoter occupancy than HA-mNTD-yTBPc, consistent with an inhibitory effect of the mouse NTD (Fig. 5D, left).

Next, the averaged reads of RNA Pol II ChIP-seq were plotted as metagene heatmaps for the 146 induced genes (Fig. 5E). Under untreated control conditions, RNA Pol II occupancy was minimal. Upon diamide treatment, all HA-tagged expressing strains showed increased RNA Pol II occupancy over the gene body from TSS to transcription end site (TES), though to different extents (Fig. 5E). For quantification, we calculated the readcounts ratio of HA-yTBPc and HA-mNTD-yTBPc over the full-length at the induced gene bodies (TSS to TES) with and without diamide (Fig. 5D, right). Both HA-yTBPc and HA-mNTD-yTBPc supported lower induction than HA-yTBP, though still above empty vector levels (Fig. 5D-E). Unexpectedly, while DNA-binding induction was stronger for HA-yTBPc than HA-mNTD-yTBPc, RNA Pol II recruitment was significantly lower for HA-yTBPc compared to the chimeric protein (Fig. 5D-E). Together, these results suggest that the TBP NTD facilitates coupling between TBP binding and RNA Pol II recruitment, thereby enhancing efficient transcriptional reprogramming under stress.

### NTD is enriched in disordered amino acids and inversely correlates with gene density across eukaryotes

The yeast and mouse NTDs differ in both length and sequence composition, yet both are highly disordered. To assess the generality of this feature across eukaryotes, we used yeast TBP (P13393) as a query in BLAST searches across five taxa (Fungi, Arthropoda, Nematoda, Vertebrates, and Mammals) recovering 402 TBP sequences from 338 species. These proteins varied in length from 153 to 394 amino acids (Fig. 6A, S6A). The taxonomic breadth was chosen to ensure representation beyond fungi. Disorder prediction scores were computed for each amino acid across the 402 sequences and visualized as a heatmap, where higher scores indicate a greater likelihood of intrinsic disorder (Fig. 6A). Alignment of the sequences at their C-terminal ends revealed consistently low disorder scores near the C-terminal region (blue), consistent with the conserved DNA-binding core domain (Fig. 6A). Representative species from each of the five taxa are highlighted (Fig. 6A, S6A). Across nearly all species, the NTDs were enriched for disordered residues, regardless of taxonomic group (Fig. 6A, S6A). The number of disordered residues positively correlated with protein length (Fig. 6B), suggesting that while the core domain has remained highly conserved, the NTD has diversified in both sequence and size throughout evolution.

**Figure 6.**
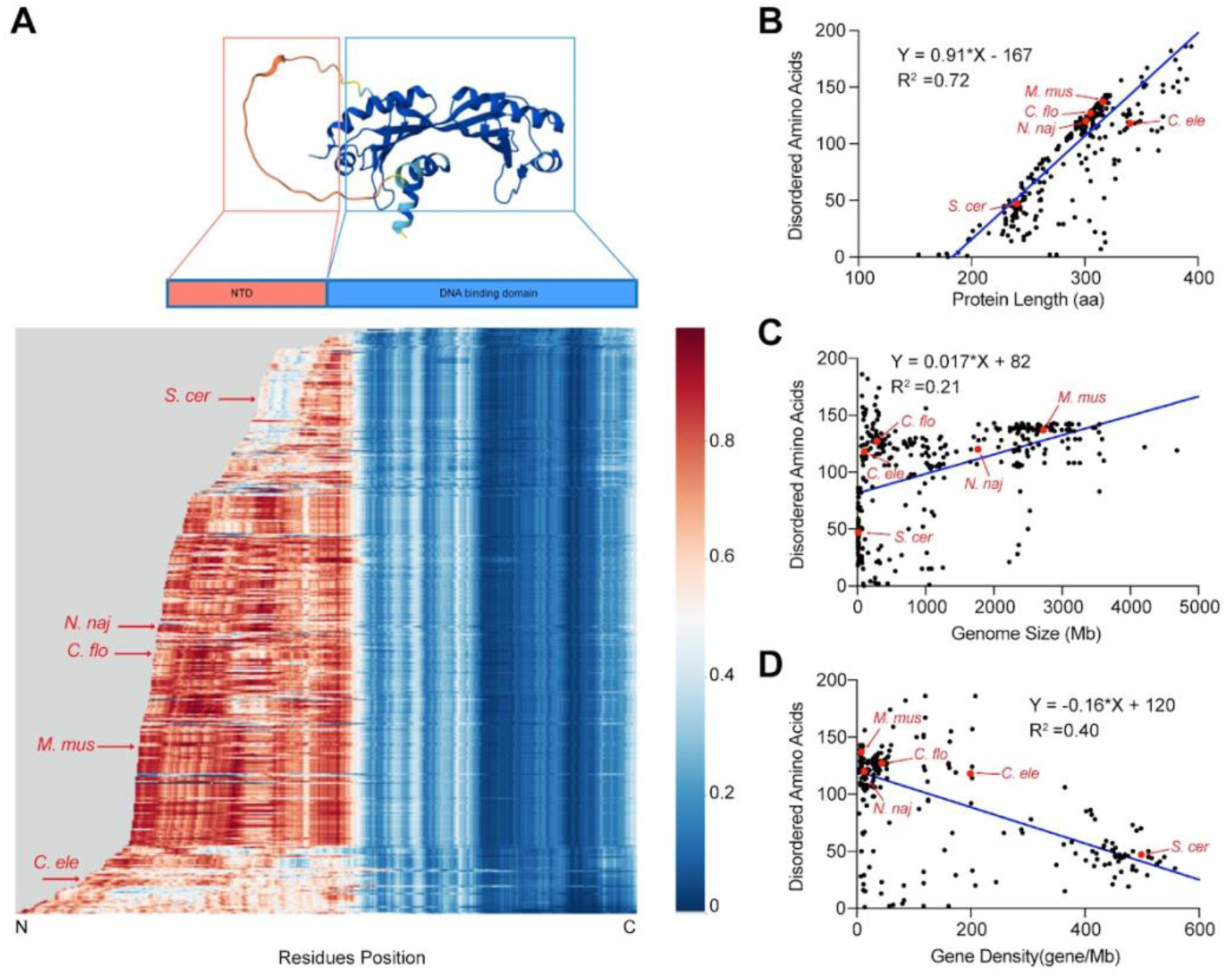
NTD is enriched in disordered amino acids, and its length inversely correlates with gene density across eukaryotes. (A) AlphaFold predicted structure of yeast TBP (P13393) (top); distribution heatmap (bottom) of disorder probability along 402 TBP sequence homologs sorted by length and aligned to C-terminal end, where a higher score represents a higher likelihood that the amino acid is in a disordered region. *S. cerevisiae (S.cer)*, *C. floridanus (C.flor)*, *C. elegans (C.ele)*, *N. naja (N.naj)*, and *M. musculus (M.mus)* were labelled as examples from five taxa (Fungi, Arthropoda, Nematoda, Vertebrates and Mammals). (B-D) The number of predicted disordered amino acids correlates positively with TBP total length (B), genome size (C) and inversely with coding gene density (D).

We next asked how TBP NTD variability relates to genome evolution, specifically genome size and gene density. Using a custom script, we compiled genome assembly sizes and annotated gene counts from the NCBI database. Gene density was calculated as total gene number divided by genome size, where lower density reflects larger non-coding regions and, by extension, greater genome complexity (Pesole *et al*, 2001; Shabalina & Spiridonov, 2004; Quintero-Cadena *et al*, 2020). The number of disordered amino acids in TBP homologs positively correlated with genome size (Fig. 6C) and inversely correlated with coding gene density (Fig. 6D) and total gene density (Fig. S6B). Taken together, these analyses suggest that organisms with more complex genomes have co-evolved longer and more disordered TBP NTDs, potentially enabling species-specific modulation of gene regulation.

## Discussion

In this study, we investigated the extent to which homologous murine TBPs can substitute for yeast TBP and examined the role of the NTD in modulating TBP activity. Using a TBP replacement assay in a temperature-sensitive yeast mutant, coupled with phenotyping, ChIP-seq, and transcriptional analyses, we found that conserved mouse TBP and its core domain partially rescue yeast viability, whereas the paralogs TRF2 and TRF3 cannot. The mouse TBP core maintained partial binding activity at RNA Pol I and II genes, but showed pronounced defects at Pol III loci. The mouse NTD exerted an inhibitory effect on both yeast and mouse TBP function, under basal and stress conditions. Finally, comparative analysis across 402 TBP homologs revealed that the NTD is universally enriched in intrinsically disordered residues, with length correlating positively with genome size and negatively with gene density. Together, these findings support a division of labor in which the conserved TBP core maintains essential promoter recognition, while the variable NTD shapes species-specific transcriptional regulation.

A central finding is that the mouse TBP core domain can partially rescue yeast TBP lethality, whereas the full-length protein performs less effectively, and paralogs fail to complement. This result underscores the deep conservation of the TBP DNA-binding core across eukaryotes, consistent with structural studies showing its stable recognition of TATA-box motifs (Kim *et al*, 1993; Ravarani *et al*, 2020). However, the incomplete rescue highlights functional divergence that cannot be explained by promoter binding alone. Previous work has demonstrated that metazoan TBPs interact with a broader repertoire of cofactors compared to yeast, and our results extend this concept by showing that the mouse NTD imposes incompatibilities when transplanted into yeast (Hampsey & Reinberg, 1999; Wu & Hampsey, 1999; Ravarani *et al*, 2020). This finding suggests that while the TBP core provides a universal scaffold for transcription initiation, the disordered domain evolves to tailor TBP function to species-specific transcriptional environments. A limitation of our study is that we examined only a single mammalian ortholog, and future work with additional TBPs from other phyla could determine whether partial rescue is a general property of all non-yeast TBPs.

Another key result is the differential ability of mouse TBP to support transcription by RNA Pol I, II, and III. While the mouse TBP core showed moderate binding at RNA Pol I promoters and retained limited function and polymerase recruitment at RNA Pol II loci, it was almost completely defective at RNA Pol III genes. This hierarchy is surprising, as TBP is considered universally required for all three RNA Pols in yeast (Cormack & Struhl, 1992b; White *et al*, 1992). Interestingly, the divergent responses we observed in yeast mirror our previous TBP swap findings in mESCs where yeast TBP was able to fully bind to RNA Pol II genes, though TBP is not strictly required in mammalian RNA Pol II transcription, but exhibited only limited binding and recruitment at Pol III genes (Kwan *et al*, 2023; Cui *et al*, 2025). Together, these results reinforce the view that RNA Pol III transcription imposes the strictest requirements on TBP–cofactor compatibility across species, whereas Pol II transcription displays greater flexibility. This divergence raises the intriguing possibility that Pol III-dependent transcription, which is tightly linked to growth and viability, enforces stricter requirements for TBP–cofactor compatibility than Pol I or II. A limitation here is that our ChIP-seq analysis may underestimate residual Pol III binding due to antibody sensitivity, but the consistent failure of mouse TBP to fully rescue Pol III function strongly supports this conclusion. Future studies could dissect which specific protein–protein interfaces between TBP and TFIIIB are disrupted by heterologous domains.

Our analysis of truncated and chimeric TBPs highlights the NTD as a critical modulator of TBP function. Although yeast NTD deletions or swaps had little impact under homeostatic conditions, they markedly impaired transcriptional induction under oxidative stress. This finding suggests that the NTD provides regulatory plasticity during transcriptional reprogramming. This result is consistent with our previous finding in mESCs that the core domain dictates the DNA-binding affinity, while the NTD has a modulatory effect, with the most effect on stress transcriptional reprogramming (Cui *et al*, 2025). Unexpectedly, we observed that the mouse NTD exerted an inhibitory effect on mouse and yeast TBP function in yeast, dampening Pol II recruitment and promoter occupancy. This finding is consistent with the notion that intrinsically disordered regions can both enable and restrict protein–protein interactions depending on context (Bondos *et al*, 2022; Ghitti *et al*, 2025; Um & Manley, 2001). These results parallel findings in other transcription factors where disordered domains confer context-dependent regulation (Csizmok *et al*, 2016; Ghitti *et al*, 2025). Future work should investigate the binding partners of the TBP NTD during stress and determine whether similar inhibitory interactions occur in mammalian systems.

Finally, our comparative analysis across 402 eukaryotic TBPs revealed that the NTD is universally enriched in intrinsically disordered residues and that its length positively correlates with genome size while inversely correlating with gene density. This analysis supports a model in which expansion of the NTD co-evolved with increasing genomic complexity, providing greater regulatory capacity to TBP in species with large, gene-sparse genomes. Such a relationship between disorder and complexity has been described for other transcriptional regulators like RNA Pol II, and our findings place TBP squarely within this paradigm (Quintero-Cadena *et al*, 2020). The implication is that the conserved TBP core maintains essential promoter recognition, while the variable NTD provides a flexible interface for regulatory evolution (Lescure *et al*, 1994; Ravarani *et al*, 2020). This raises new questions about how disordered domains expand the functional repertoire of otherwise highly conserved transcription factors and whether NTD variability across species corresponds to differences in transcriptional network architecture.

## Materials and Methods

### Reagents and Tools Table

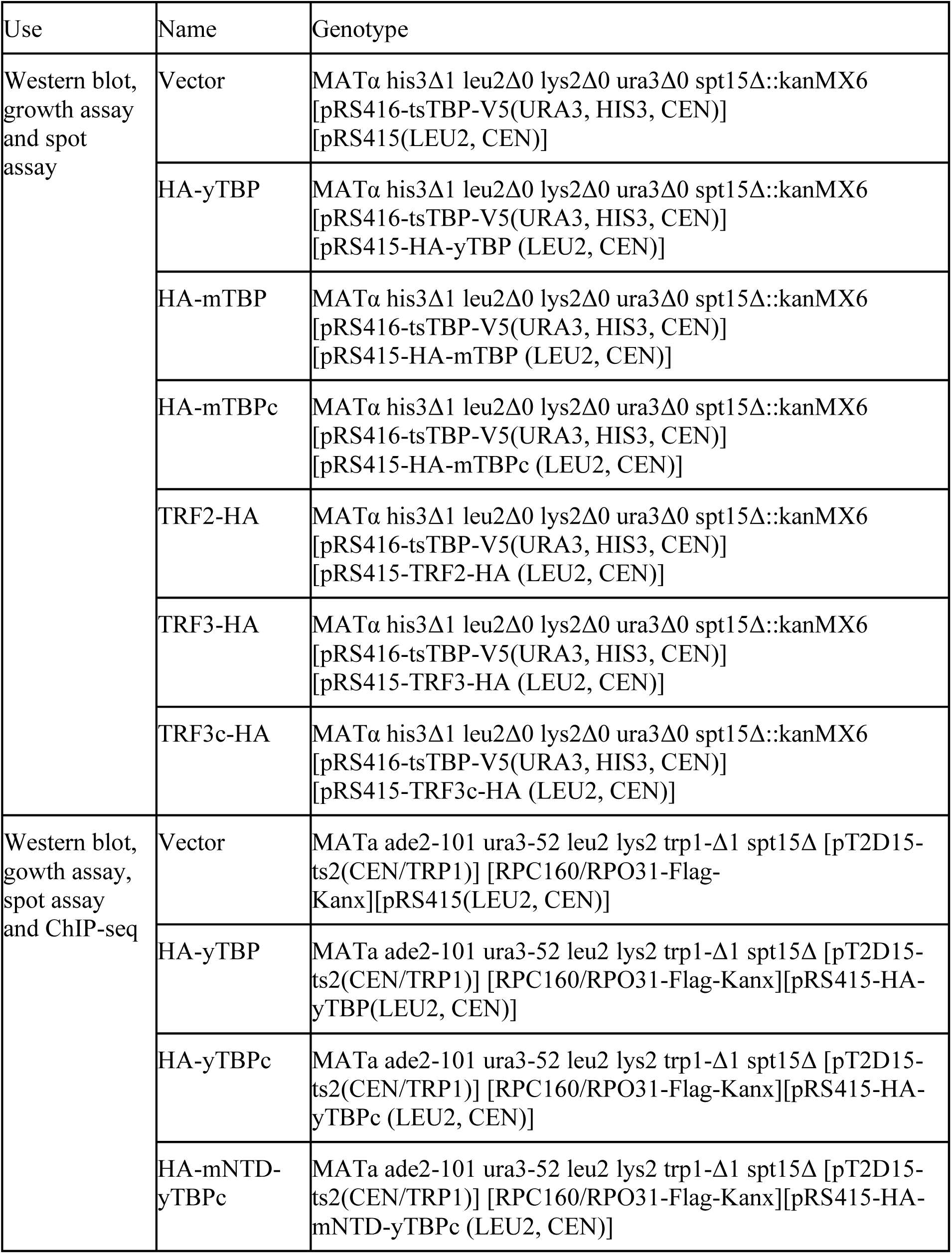
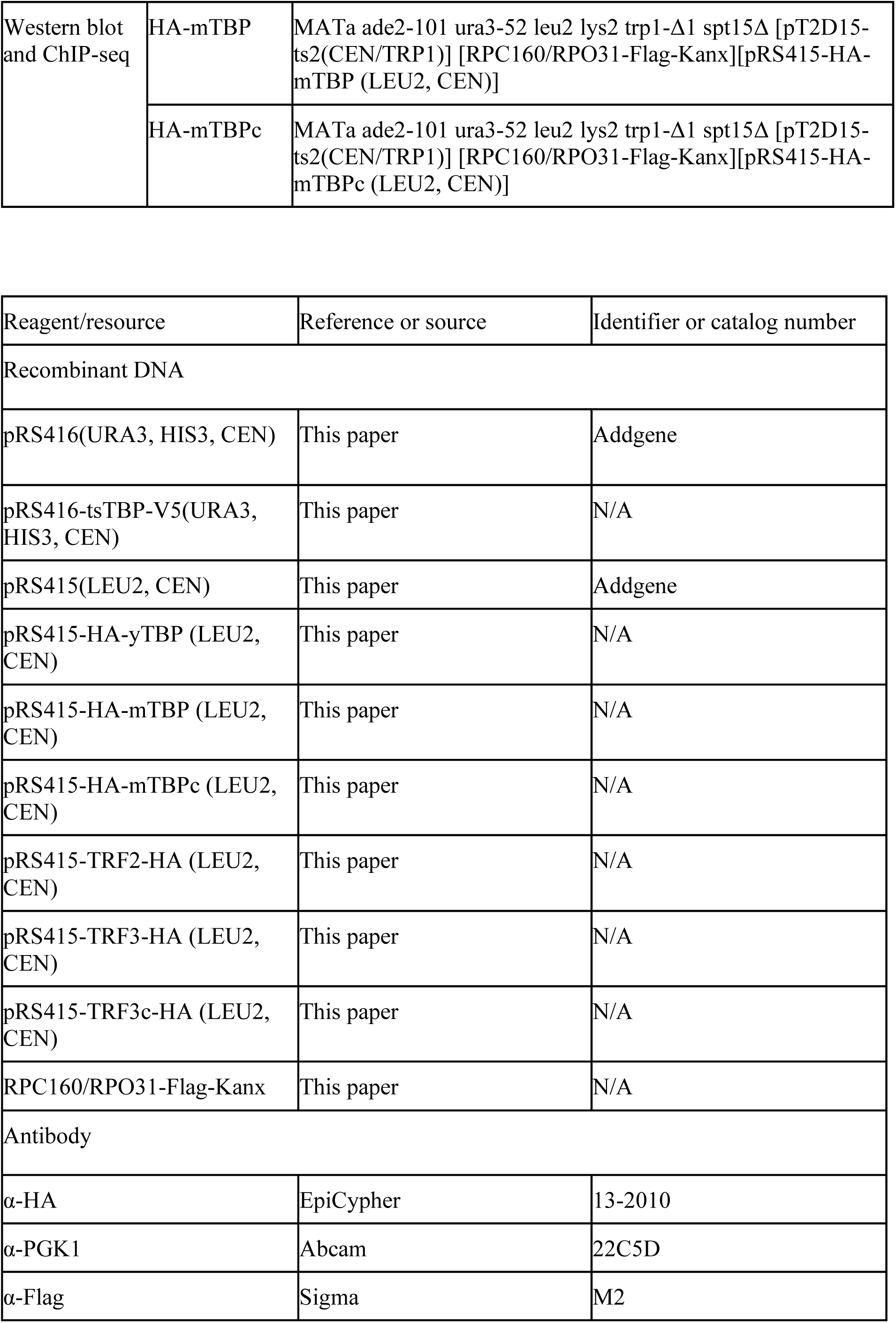
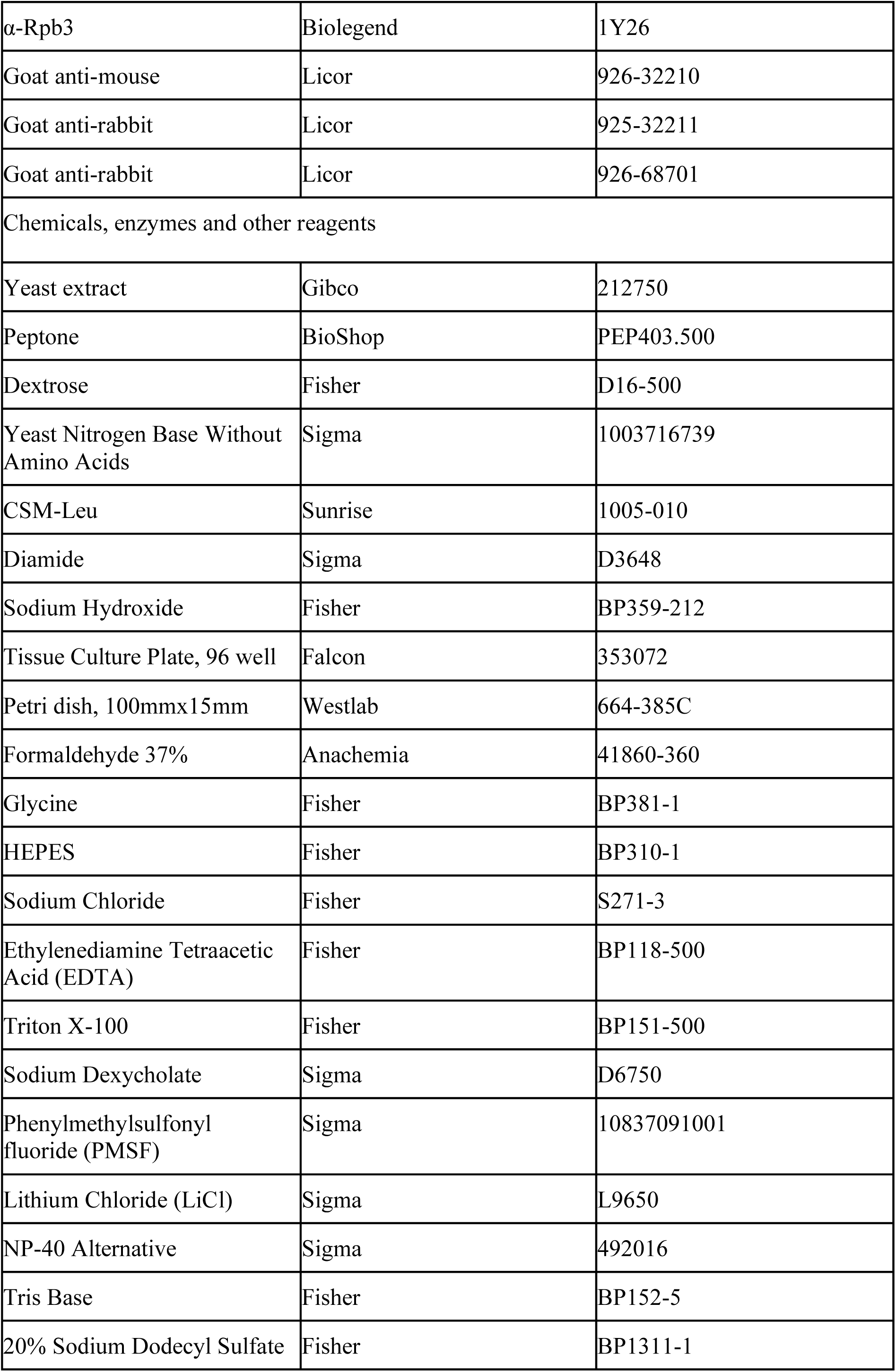
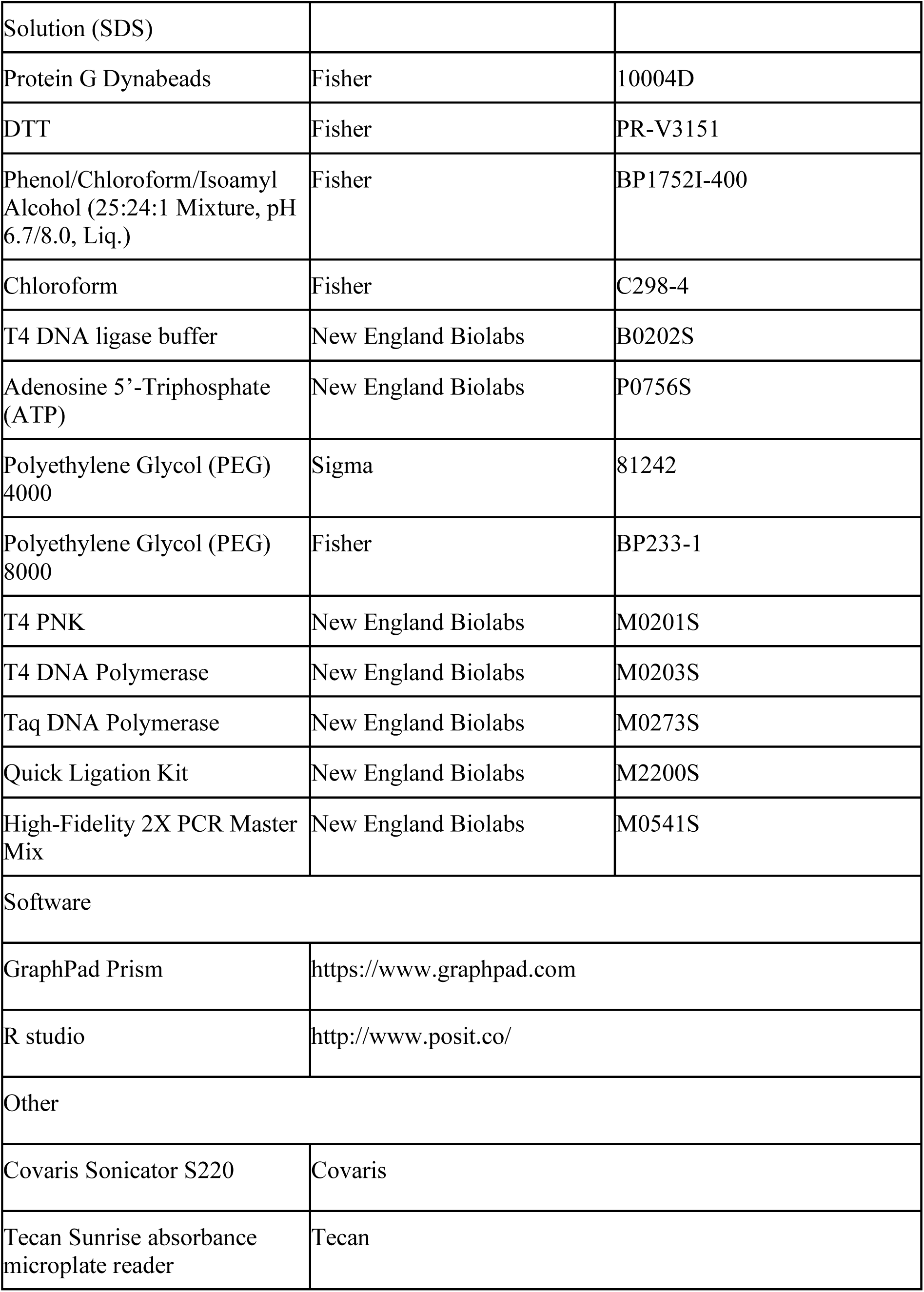

## Methods and Protocols

### Yeast strain and growth

All strains used in this study were isogenic to BY2, purchased from NBRP (National BioResource Project) and are listed in Reagents and Tools Table. Yeast cultures were grown in either selective media after transformation or YPD media (1% yeast extract, 2% peptone, and 2% dextrose). Diamide treatments were performed in YPD with 0.75mM diamide for 15 minutes, unless noted otherwise.

### RPC160 Knock in

The knock in was performed as previously described (Funakoshi & Hochstrasser, 2009) with primer sequence, forward: ATTTGAAAGTCTCTCAAATGAGGCAGCTTTAAAAGCGAACgactacaaagaccatgacg; reverse: ATACAAATGCTATAAAAAAGTTTAAAAACGACTACTTTAcagtatagcgaccagcattc.

### Whole cell extraction and antibodies used for Western blot analyses

Whole-cell extracts were generated as previously described (Kushnirov, 2000). Primary antibodies: anti-HA (α-HA) 1:5000 (EpiCypher 13-2010), α-PGK1 1:5000 (Abcam 22C5D8), α-Flag 1:5000 (Sigma M2). Secondary antibodies: IRDye 800CW Goat anti-mouse (Licor 926-32210), IRDye 800CW Goat anti-rabbit (Licor 925-32211), IRDye 680RD Goat anti-mouse (Licor 926-68070), or IRDye 680RD Goat anti-rabbit (Licor 926-68701).

### Spot assay

Cells at the exponential phase were collected and washed with sdH2O, and serially diluted on YPD plates. Plates were incubated at the permissive (29 ± 0.5 °C) or restrictive temperature (36.5 ± 0.5 °C)for 24 to 36 hours.

### Growth assay

Optical density (OD) was measured at an absorbance wavelength of 620 nm, for 0 to 24 hours every 5 minutes. 1:100 dilutions of overnight cultures were grown in 100 uL of YPD, in a Falcon flat-bottom 96-Well Clear Assay Plate with lid, on Tecan microplate reader with shaking at both the permissive (29 ± 0.5 °C) and restrictive temperature (36.5 ± 0.5 °C) as previously described (Toussaint & Conconi, 2006). The growth curves were quantified through the area under the fitted logistic curve using Growthcurver.

### Chromatin immunoprecipitation sequencing

Cells were crosslinked in 1% formaldehyde for 20 minutes and quenched with the addition of liquid glycine to 300 mM for a further 5 minutes at room temperature. Cells were lysed by bead beating, and cell lysate was spun down at 15,000g for 30 minutes. The cell pellet was resuspended in lysis buffer (50 mM HEPES, pH 7.5, 140 mM NaCl, 1 mM EDTA, 1% Triton X-100, 0.1% sodium deoxycholate) and sonicated (Covaris Sonicator S220) to produce an average fragment size of 250 bp. The lysate was spun down at 9000g for 10 minutes, and the supernatant was precleared by rotating with Protein G Dynabeads for 1 hour at 4 °C. Five percent of the lysate was reserved for input, and the remaining was split and incubated with α-HA (EpiCypher 13-2010), α-Rpb3 (Biolegend 1Y26) or α-Flag (Sigma M2) antibodies overnight at 4 °C, respectively.

Antibody immunoprecipitations were isolated by adding magnetic Protein G Dynabeads and rotating at 4 °C for 1 h, and 5 minute washes were performed twice with lysis buffer, twice with high salt buffer (50 mM HEPES pH 7.5, 500 mM NaCl, 1 mM EDTA, 1% Triton X-100, 0.1% sodium-deoxycholate), twice with LiCl wash buffer (10 mM Tris-HCl pH 8.0, 250 mM LiCl, 0.6% NP-40, 0.5% sodium-deoxycholate, 1 mM EDTA), and once with TE (10mM Tris pH 8.0, 1 mM EDTA). Synthetic spike-in *E coli* DNA was added to eluates, to aid in quantification. Following proteinase K digestion, DNA was purified by phenol, chloroform, isoamyl alcohol extraction and RNase A treated.

### Library preparation

ChIP-seq libraries were prepared using the the Henikoff lab in house library prep strategy for Illumina sequencing that uses TruSeq-Y Adapters with a free 3’T overhang (Janssens & Henikoff, 2019).

### Data analysis

Reads were trimmed for adapters sequence GATCGGAAGAGCACACGTCTGAACTCCAGTCA and mapped on sacCre3 genome build using Bowtie2 with the following parameters: --local --very-sensitive-local --no-unal --no-mixed --no-discordant --phred33 -I 10 -X 700. Polymerase chain reaction (PCR) duplicate reads were removed. A normalization factor was determined from *E coli* DNA (normalized to the control) alignment from Bowtie2 mapping and used to scale during the generation of bigwig files. Replicates were averaged using bigwigAverage, which averages the reads after normalization. Downstream analyses, heatmaps, TSS plots, gene plots, and k-means clustering were performed using IGV, DeepTools, and BedTools suite. ComputeMatrix from deeptools was done using binsize 10.

### Construction of a customized yeast genome for ribosomal DNA mapping

The Saccharomyces cerevisiae reference genome R64-1-1 and its annotations were downloaded from Ensembl release 111 (Dyer *et al*, 2025). The coordinates and annotations labelled with the biotype rRNA were extracted from the reference gff3 annotation file and the corresponding sequences in the reference genome were masked using the BEDTools suite after being extended by 100 bases on both sides. The original sequence of length 9135 starting at coordinate 459797 on chromosome XII was then added to the customized genome with the label rDNA. A few additional regions with significant blastn hits to that rDNA sequence were also masked. The annotations of the rRNA genes were then modified to take into account the new sequence label and coordinates.

### TBP bioinformatic analysis

TBP homologs were retrieved by searching the Unpriot with default settings, starting with the yeast TBP protein sequence (G4XSG8), in Fungi, Arthropoda, Nematoda, Vertebrate and Mammal for the top 100 hits. The results were sorted and cleaned up by removing duplicated protein entries. 402 amino acid sequences were analyzed for disorder using Iupred2A with default settings, which provides a disordered score for each amino acid (Mészáros *et al*, 2018). Genome sizes and gene numbers were scraped from NCBI websites using a custom script (Chan, 2024).

### Data, code, and materials availability

All sequencing data have been deposited in Gene Expression Omnibus (Accession number: GSE308766). All other data are available in the manuscript or in the supplementary materials.

### Supplementary Data

Supplementary table_1_TBP_length

Supplementary table_2_total_gene_density

Supplementary table_3_coding_gene_density

## Acknowledgments

We thank Dr. Eric Jan for providing S2 cells for spike-in normalization. We thank T. Stach (BRC-seq, UBC) for Illumina sequencing. This work was supported by Life Sciences Institute Cores (LSI Imaging, ubcFLOW, and QPCR Core), and by the UBC GREx Biological Resilience Initiative. For insightful comments on the manuscript, we thank Dr. Leann Howe. J.C. is supported by UBC 4-Year Fellowship, S.S.T is a Canada Research Chair and Michael Smith Foundation for Health Research Scholar

## Author contribution

Conceptualization: JHC, SST

Methodology: JHC, HY, DC, SF

Investigation: JHC, HY, DC

Visualization: JHC, SST

Funding acquisition: JHC, SST

Supervision: SST

Writing – original draft: JHC, SST

Writing – review and editing: JHC, SST

## Reference

1. Akhtar W & Veenstra GJC (2009) TBP2 is a substitute for TBP in Xenopus oocyte transcription. BMC Biol 7: 45

2. Akhtar W & Veenstra GJC (2011) TBP-related factors: a paradigm of diversity in transcription initiation. Cell Biosci 1: 23

3. Bondos SE, Dunker AK & Uversky VN (2022) Intrinsically disordered proteins play diverse roles in cell signaling. Cell Commun Signal 20: 20

4. Buratowski S, Hahn S, Sharp PA & Guarente L (1988) Function of a yeast TATA element-binding protein in a mammalian transcription system. Nature 334: 37–42

5. Chan FC (2024) Get the reference assembly length and number of protein coding genes per taxon.

6. Cormack BP, Strubin M, Stargell LA & Struhl K (1994) Conserved and nonconserved functions of the yeast and human TATA-binding proteins. Genes Dev 8: 1335–1343

7. Cormack BP & Struhl K (1992a) The TATA-binding protein is required for transcription by all three nuclear RNA polymerases in yeast cells. Cell 69: 685–696

8. Cormack BP & Struhl K (1992b) The TATA-binding protein is required for transcription by all three nuclear RNA polymerases in yeast cells. Cell 69: 685–696

9. Csizmok V, Follis AV, Kriwacki RW & Forman-Kay JD (2016) Dynamic Protein Interaction Networks and New Structural Paradigms in Signaling. Chem Rev 116: 6424–6462

10. Cui JH, Kwan JZJ, Faghihi A, Nguyen TF & Teves SS (2025) Functional divergence of TBP homologs through distinct DNA-binding dynamics. Nucleic Acids Res 53: gkaf436

11. Dieci G, Conti A, Pagano A & Carnevali D (2013) Identification of RNA polymerase III-transcribed genes in eukaryotic genomes. Biochim Biophys Acta BBA - Gene Regul Mech 1829: 296–305

12. Dyer SC, Austine-Orimoloye O, Azov AG, Barba M, Barnes I, Barrera-Enriquez VP, Becker A, Bennett R, Beracochea M, Berry A, et al (2025) Ensembl 2025. Nucleic Acids Res 53: D948–D957

13. French SL, Osheim YN, Schneider DA, Sikes ML, Fernandez CF, Copela LA, Misra VA, Nomura M, Wolin SL & Beyer AL (2008) Visual Analysis of the Yeast 5S rRNA Gene Transcriptome: Regulation and Role of La Protein. Mol Cell Biol 28: 4576–4587

14. Funakoshi M & Hochstrasser M (2009) Small epitope-linker modules for PCR-based C-terminal tagging in Saccharomyces cerevisiae. Yeast Chichester Engl 26: 185–192

15. Gasch AP, Spellman PT, Kao CM, Carmel-Harel O, Eisen MB, Storz G, Botstein D & Brown PO (2000) Genomic Expression Programs in the Response of Yeast Cells to Environmental Changes. Mol Biol Cell 11: 4241–4257

16. George SS, Pimkin M & Paralkar VR (2023) Construction and validation of customized genomes for human and mouse ribosomal DNA mapping. J Biol Chem 299: 104766

17. Ghitti M, Colley LS, Mantonico MV, Musco G & Bianchi ME (2025) Intrinsic disorder and fuzzy interactions drive multiple functions of HMGB1. Trends Biochem Sci: S0968000425001902

18. Hampsey M & Reinberg D (1999) RNA polymerase II as a control panel for multiple coactivator complexes. Curr Opin Genet Dev 9: 132–139

19. Hobbs NK, Bondareva AA, Barnett S, Capecchi MR & Schmidt EE (2002) Removing the Vertebrate-Specific TBP N Terminus Disrupts Placental β2m-Dependent Interactions with the Maternal Immune System. Cell 110: 43–54

20. Hoey T, Dynlacht BD, Peterson MG, Pugh BF & Tjian R (1990) Isolation and characterization of the Drosophila gene encoding the TATA box binding protein, TFIID. Cell 61: 1179–1186

21. Hurowitz EH & Brown PO (2003) Genome-wide analysis of mRNA lengths in Saccharomyces cerevisiae. Genome Biol 5: R2

22. Janssens D & Henikoff S (2019) CUT&RUN: Targeted in situ genome-wide profiling with high efficiency for low cell numbers v3. doi:10.17504/protocols.io.zcpf2vn [PREPRINT]

23. Kim JL, Nikolov DB & Burley SK (1993) Co-crystal structure of TBP recognizing the minor groove of a TATA element. Nature 365: 520–527

24. Kim Y-H (2006) Chromosome XII context is important for rDNA function in yeast. Nucleic Acids Res 34: 2914–2924

25. Kushnirov VV (2000) Rapid and reliable protein extraction from yeast. Yeast 16: 857–860

26. Kwan JZ, Nguyen TF & Teves SS (2024) TBP facilitates RNA Polymerase I transcription following mitosis. RNA Biol 21: 42

27. Kwan JZ, Nguyen TF, Uzozie AC, Budzynski MA, Cui J, Lee JM, Van Petegem F, Lange PF & Teves SS (2023) RNA Polymerase II transcription independent of TBP in murine embryonic stem cells. eLife 12: e83810

28. Kwon H & Green MR (1994) The RNA polymerase I transcription factor, upstream binding factor, interacts directly with the TATA box-binding protein. J Biol Chem 269: 30140–30146

29. Lee DK, Horikoshi M & Roeder RG (1991) Interaction of TFIID in the minor groove of the TATA element. Cell 67: 1241–1250

30. Lee M & Struhl K (2001) Multiple functions of the nonconserved N-terminal domain of yeast TATA-binding protein. Genetics 158: 87–93

31. Lescure A, Lutz Y, Eberhard D, Jacq X, Krol A, Grummt I, Davidson I, Chambon P & Tora L (1994) The N-terminal domain of the human TATA-binding protein plays a role in transcription from TATA-containing RNA polymerase II and III promoters. EMBO J 13: 1166–1175

32. Martianov I, Fimia GM, Dierich A, Parvinen M, Sassone-Corsi P & Davidson I (2001) Late arrest of spermiogenesis and germ cell apoptosis in mice lacking the TBP-like TLF/TRF2 gene. Mol Cell 7: 509–515

33. Matsui T, Segall J, Weil PA & Roeder RG (1980) Multiple factors required for accurate initiation of transcription by purified RNA polymerase II. J Biol Chem 255: 11992–11996

34. Mészáros B, Erdős G & Dosztányi Z (2018) IUPred2A: context-dependent prediction of protein disorder as a function of redox state and protein binding. Nucleic Acids Res 46: W329– W337

35. Nakajima N, Horikoshi M & Roeder RG (1988) Factors Involved in Specific Transcription by Mammalian RNA Polymerase II: Purification, Genetic Specificity, and TATA Box-Promoter Interactions of TFIID. Mol Cell Biol 8: 4028–4040

36. Paule MR (2000) SURVEY AND SUMMARY Transcription by RNA polymerases I and III. Nucleic Acids Res 28: 1283–1298

37. Pesole G, Mignone F, Gissi C, Grillo G, Licciulli F & Liuni S (2001) Structural and functional features of eukaryotic mRNA untranslated regions. Gene 276: 73–81

38. Petes TD (1979) Yeast ribosomal DNA genes are located on chromosome XII. Proc Natl Acad Sci 76: 410–414

39. Petrenko N, Jin Y, Dong L, Wong KH & Struhl K (2019) Requirements for RNA polymerase II preinitiation complex formation in vivo. eLife 8: e43654

40. Petrenko N, Jin Y, Wong KH & Struhl K (2017) Evidence that Mediator is essential for Pol II transcription, but is not a required component of the preinitiation complex in vivo. eLife 6: e28447

41. Quintero-Cadena P, Lenstra TL & Sternberg PW (2020) RNA Pol II Length and Disorder Enable Cooperative Scaling of Transcriptional Bursting. Mol Cell 79: 207–220.e8

42. Ravarani CNJ, Flock T, Chavali S, Anandapadamanaban M, Babu MM & Balaji S (2020) Molecular determinants underlying functional innovations of TBP and their impact on transcription initiation. Nat Commun 11: 2384

43. Roeder RG & Rutter WJ (1969) Multiple Forms of DNA-dependent RNA Polymerase in Eukaryotic Organisms. Nature 224: 234–237

44. Rowlands T, Baumann P & Jackson SP (1994) The TATA-Binding Protein: a General Transcription Factor in Eukaryotes and Archaebacteria. Science 264: 1326–1329

45. Santana JF, Collins GS, Parida M, Luse DS & Price DH (2022) Differential dependencies of human RNA polymerase II promoters on TBP, TAF1, TFIIB and XPB. Nucleic Acids Res 50: 9127–9148

46. Schramm L & Hernandez N (2002) Recruitment of RNA polymerase III to its target promoters. Genes Dev 16: 2593–2620

47. Schröder O, Bryant GO, Geiduschek EP, Berk AJ & Kassavetis GA (2003) A common site on TBP for transcription by RNA polymerases II and III. EMBO J 22: 5115–5124

48. Segall J, Matsui T & Roeder RG (1980) Multiple factors are required for the accurate transcription of purified genes by RNA polymerase III. J Biol Chem 255: 11986–11991

49. Shabalina SA & Spiridonov NA (2004) The mammalian transcriptome and the function of non-coding DNA sequences. Genome Biol 5: 105

50. Shen Y, Kassavetis GA, Bryant GO & Berk AJ (1998) Polymerase (Pol) III TATA Box-Binding Protein (TBP)-Associated Factor Brf Binds to a Surface on TBP Also Required for Activated Pol II Transcription. Mol Cell Biol 18: 1692–1700

51. Starr DB & Hawley DK (1991) TFIID binds in the minor groove of the TATA box. Cell 67: 1231–1240

52. Steffan JS, Keys DA, Dodd JA & Nomura M (1996) The role of TBP in rDNA transcription by RNA polymerase I in Saccharomyces cerevisiae: TBP is required for upstream activation factor-dependent recruitment of core factor. Genes Dev 10: 2551–2563

53. Takada S, Lis JT, Zhou S & Tjian R (2000) A TRF1:BRF Complex Directs Drosophila RNA Polymerase III Transcription. Cell 101: 459–469

54. Toussaint M & Conconi A (2006) High-throughput and sensitive assay to measure yeast cell growth: a bench protocol for testing genotoxic agents. Nat Protoc 1: 1922–1928

55. Um M & Manley JL (2001) The Drosophila TATA Binding Protein Contains a Strong But Masked Activation Domain. Gene Expr 9: 123–132

56. Werner F & Grohmann D (2011) Evolution of multisubunit RNA polymerases in the three domains of life. Nat Rev Microbiol 9: 85–98

57. White RJ, Rigby PWJ & Jackson SP (1992) The TATA-binding protein is a general transcription factor for RNA polymerase III. J Cell Sci 1992: 1–7

58. Wu W-H & Hampsey M (1999) Transcription: Common cofactors and cooperative recruitment. Curr Biol 9: R606–R609

59. Yu C, Cvetesic N, Hisler V, Gupta K, Ye T, Gazdag E, Negroni L, Hajkova P, Berger I, Lenhard B, et al (2020) TBPL2/TFIIA complex establishes the maternal transcriptome through oocyte-specific promoter usage. Nat Commun 11: 6439

60. Zhang D, Penttila T-L, Morris PL & Roeder RG (2001) Cell- and stage-specific high-level expression of TBP-related factor 2 (TRF2) during mouse spermatogenesis. Mech Dev 106: 203–205

