## Supplementary Figures and Legend for "Molecular determinants underlying functional divergence of TBP homologs"

### Supplementary Figure Legends

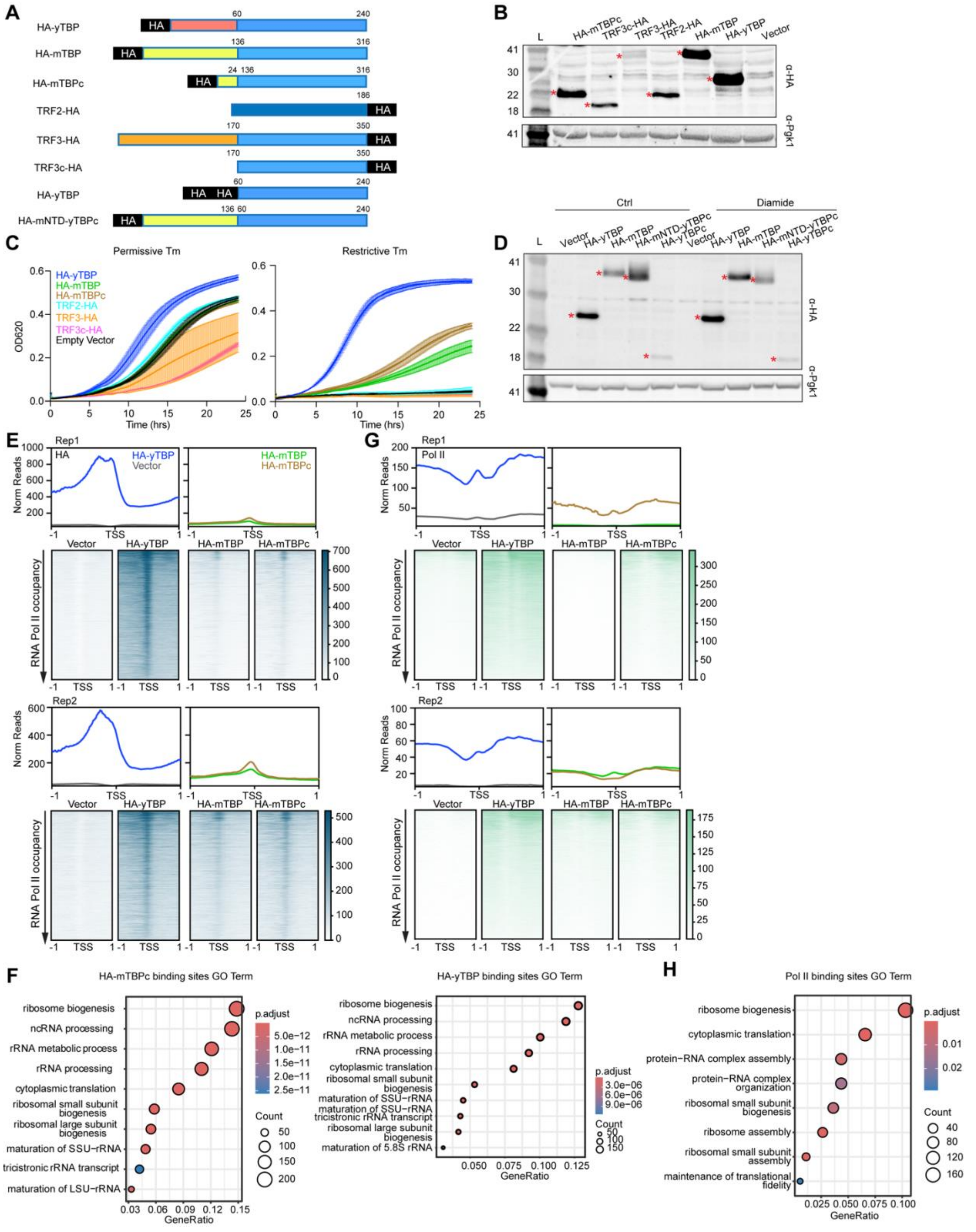

**Figure S1. TBP homologs expression and replicate analyses of TBP homologs' DNA binding at all genes.**

(A) Depiction of each HA-tagged construct aligned at the core domain are shown. (B) Western blot analyses of each HA-tagged homolog are shown. Loading controls are depicted using α-Pgk1. Asterisk for protein location. (C) Growth assay of the parental ts mutant expressing HA-yTBP, HA-mTBP, HA-mTBPc, TRF2-HA, TRF3-HA, TRF3c-HA and vector at the permissive temperature (left) and at the restrictive temperature (right). (D) Western blot analyses of each HA-tagged homolog collected at the restrictive temperature with (Diamide) and without (Ctrl) diamide are shown. Loading controls are depicted using α-Pgk1. Asterisk for protein location. (E) Biological replicates of HA ChIP-seq for HA-yTBP and vector (left), HA-mTBP and HA-mTBPc, arranged by decreasing RNA Pol II occupancy, displayed as average plots (top) and heatmaps (bottom) in a 2-kb window centered at the TSS of all genes. (F) GO term identified for HA-mTBPc and HA-yTBP significantly bound sites. (G) Biological replicates of RNA Pol II ChIP-seq for HA-yTBP and vector (left), HA-mTBP and HA-mTBPc, arranged by decreasing RNA Pol II occupancy, displayed as average plots (top) and heatmaps (bottom) in a 2-kb window centered at the TSS of all genes. (H) GO term identified for RNA Pol II bound sites in HA-mTBPc strain over vector strain. Statistics calculated using one-way ANOVA: *p-value $\leq 0.1$, **p-value $\leq 0.01$,*** p-value $\leq 0.001$; Non-significant is not shown.

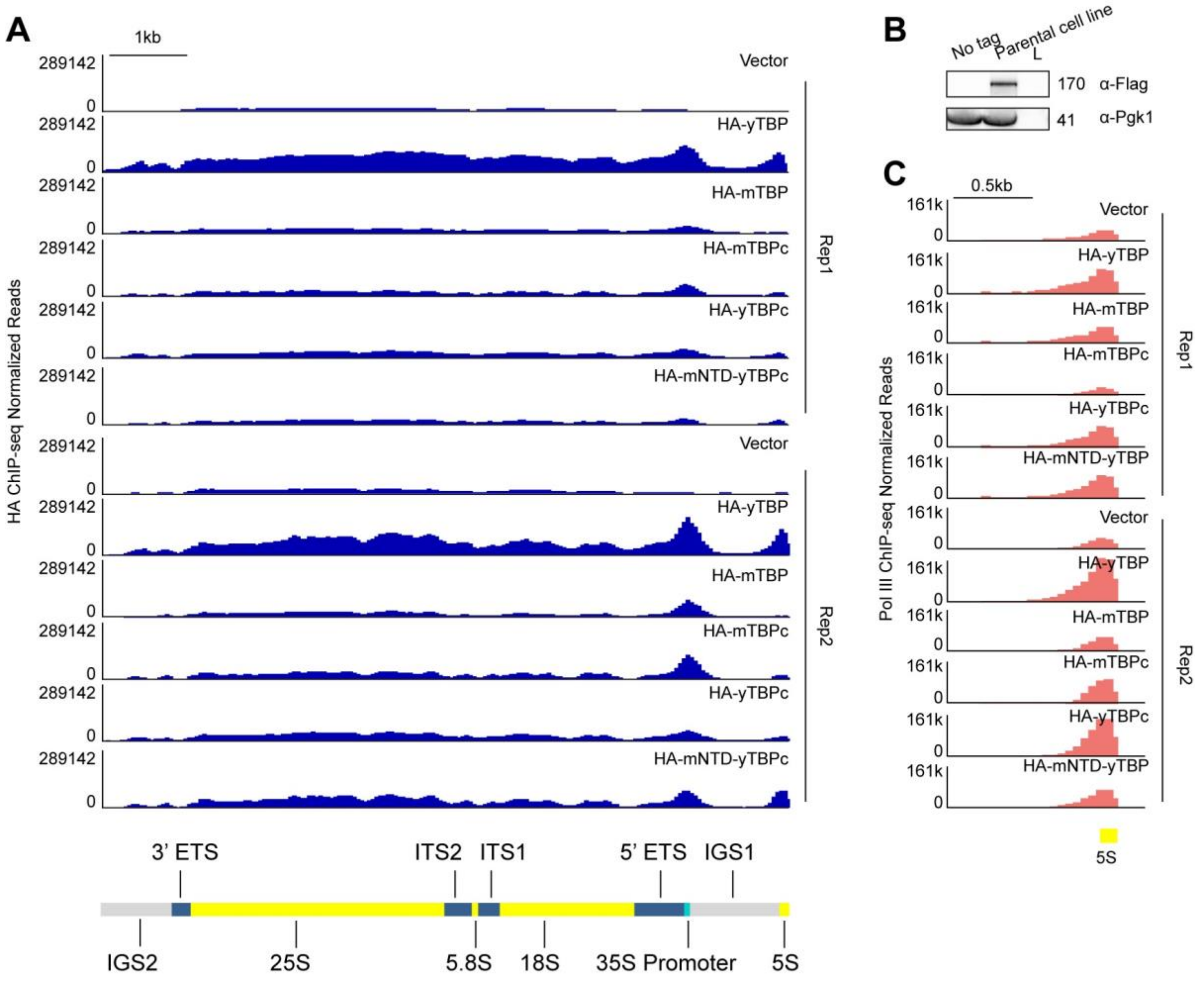


**Figure S2. Replicate analyses of TBP homologs' DNA binding at the customized rDNA chromosome.**

(A) Gene browser tracks of the custom rDNA genome on the entire rDNA chromosome (chrR) for each biological replicate of HA ChIP-seq normalized reads of vector, HA-yTBP, HA-mTBP, HA-mTBPc, HA-yTBPc and HA-mNTD-yTBPc. (B) Western blot analyses of the Flag knock-in to the parental cell line. Loading controls are depicted using α-Pgk1. (C) Gene browser tracks of the 5S rDNA for each biological replicate of HA ChIP-seq normalized reads of vector, HA-yTBP, HA-mTBP, HA-mTBPc, HA-yTBPc and HA-mNTD-yTBPc.

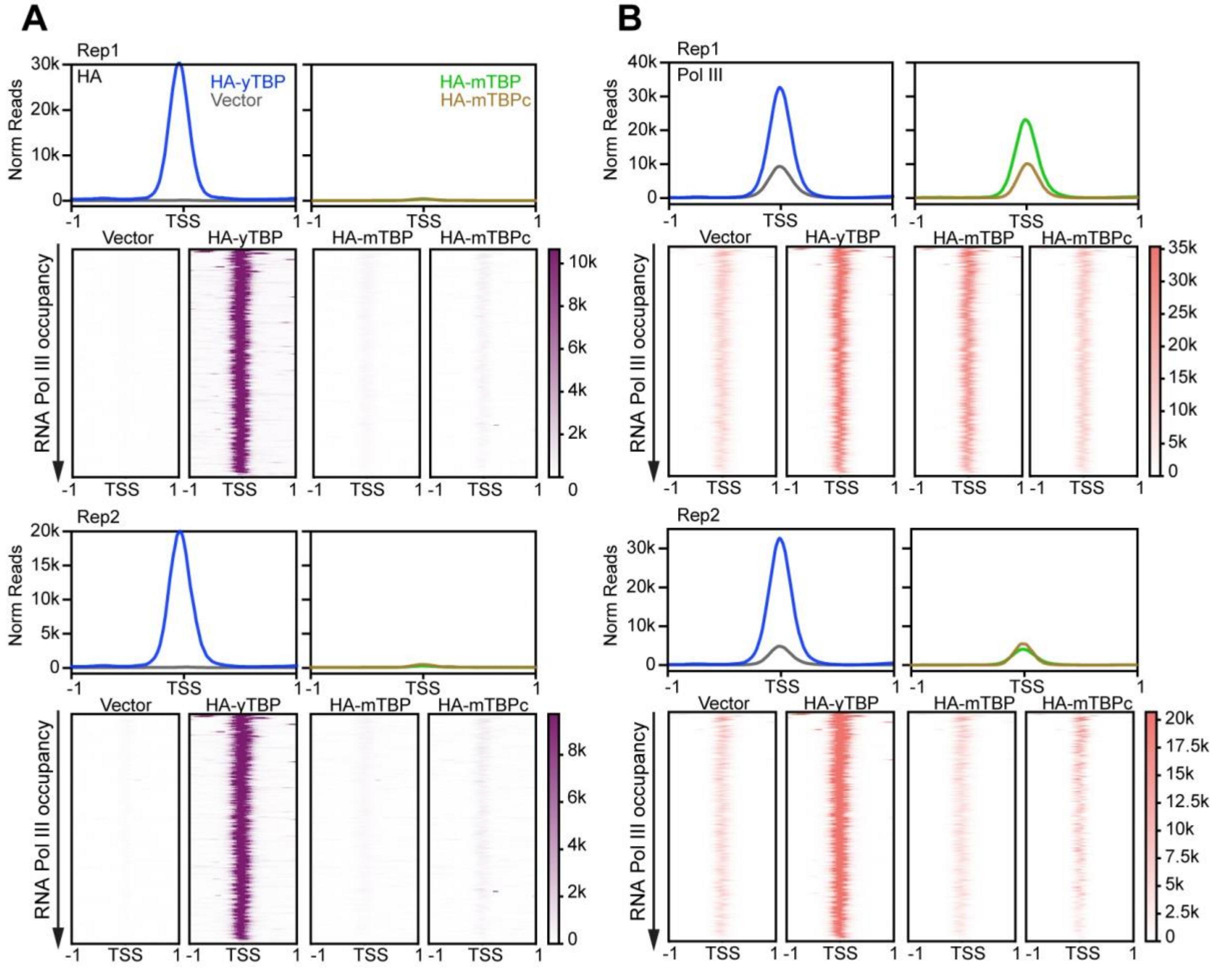


**Figure S3. Replicate analyses of TBP homologs' DNA binding at the tRNA genes.**

(A) Biological replicates of HA ChIP-seq for HA-yTBP and vector (left), HA-mTBP and HA-mTBPc, arranged by decreasing RNA Pol III occupancy, displayed as average plots (top) and heatmaps (bottom) in a 2-kb window centered at the TSS of all tRNA genes. (B) Biological replicates of RNA Pol III ChIP-seq for HA-yTBP and vector (left), HA-mTBP and HA-mTBPc, arranged by decreasing RNA Pol III occupancy, displayed as average plots (top) and heatmaps (bottom) in a 2-kb window centered at the TSS of all tRNA genes.

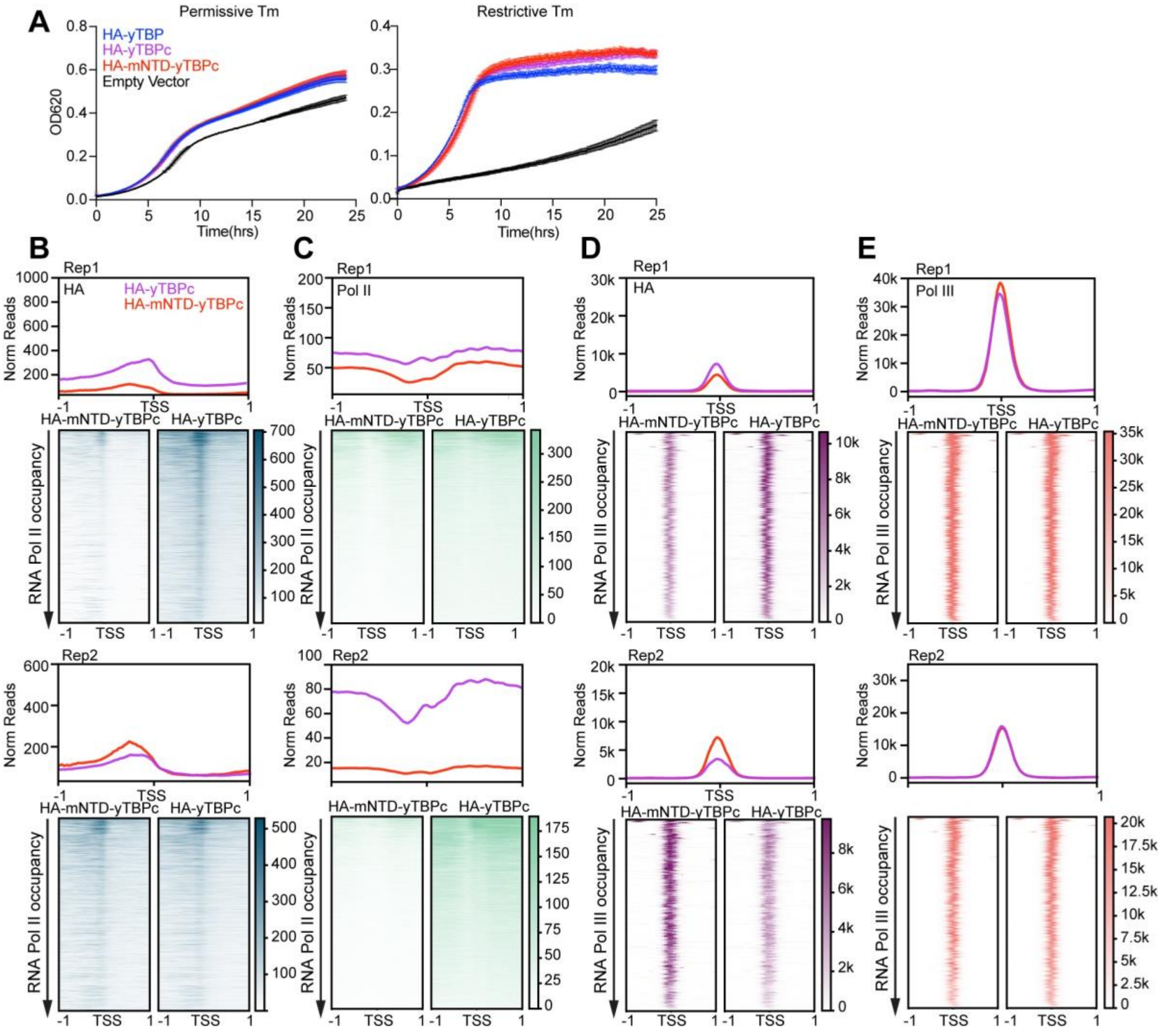


**Figure S4. Replicate analyses of TBP truncation and chimeric constructs' DNA binding activity at hemostasis.**

(A) Growth assay of parental ts mutant expressing HA-yTBP, HA-yTBPc, HA-mNTD-yTBPc and vector at the permissive temperature (left) and at the restrictive temperature (right). (B) Biological replicates of HA ChIP-seq for HA-yTBPc and HA-mNTD-yTBPc, arranged by decreasing RNA Pol II occupancy, displayed as average plots (top) and heatmaps (bottom) in a 2-kb window centered at the TSS of all genes. (C) Biological replicates of RNA Pol II ChIP-seq for HA-yTBPc and HA-mNTD-yTBPc, arranged by decreasing RNA Pol II occupancy, displayed as average plots (top) and heatmaps (bottom) in a 2-kb window centered at the TSS of all genes. (D) Biological replicates of HA ChIP-seq for HA-yTBPc and HA-mNTD-yTBPc, arranged by decreasing RNA Pol III occupancy, displayed as average plots (top) and heatmaps (bottom) in a 2-kb window centered at all the tRNA genes. (E) Biological replicates of RNA Pol III ChIP-seq for HA-yTBPc and HA-mNTD-yTBPc, arranged by decreasing RNA Pol III occupancy, displayed as average plots (top) and heatmaps (bottom) in a 2-kb window centered at all the tRNA genes. Statistics calculated using one-way ANOVA: *** p-value ≤ 0.001; Non-significant is not shown.

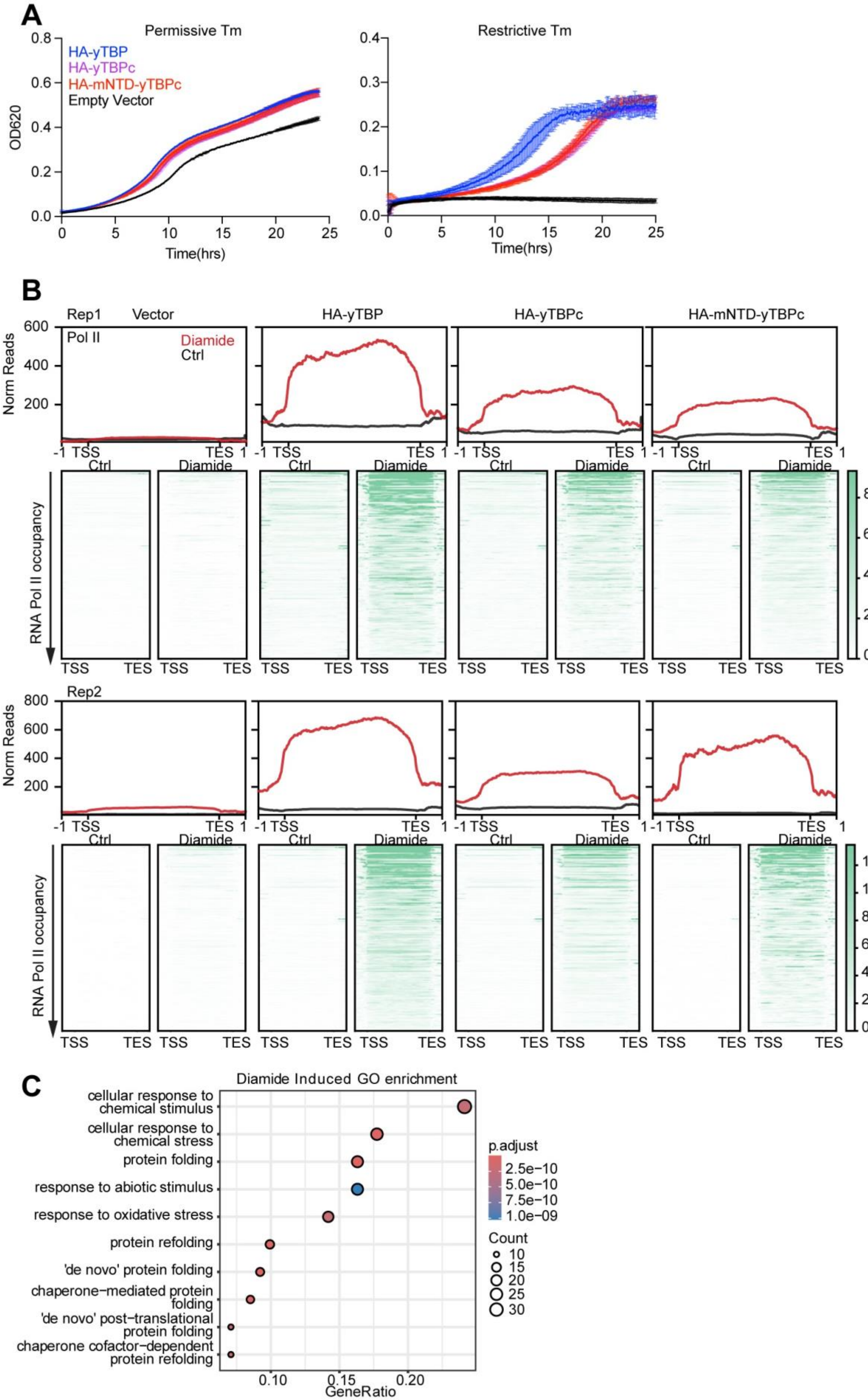
A
Permissive Tm
Restrictive Tm
HA-yTBP
HA-yTBPc
HA-mNTD-yTBPc
Empty Vector
OD620
Time(hrs)
B
Rep1
Vector
HA-yTBP
HA-yTBPc
HA-mNTD-yTBPc
Pol II
Diamide
Ctrl
Norm Reads
-1 TSS
TES 1
RNA Pol II occupancy
TSS
TES
Rep2
C
Diamide Induced GO enrichment
cellular response to chemical stimulus
cellular response to chemical stress
protein folding
response to abiotic stimulus
response to oxidative stress
protein refolding
'de novo' protein folding
chaperone−mediated protein folding
'de novo' post−translational protein folding
chaperone cofactor−dependent protein refolding
p.adjust
2.5e−10
5.0e−10
7.5e−10
1.0e−09
Count
10
15
20
25
30
GeneRatio

**Figure S5. Replicate analyses of TBP truncation and chimeric constructs' DNA binding activity at diamide-induced genes.**

(A) Growth assay of the parental ts mutant expressing HA-yTBP, HA-yTBPc, HA-mNTD-yTBPc and vector at the permissive temperature with diamide (left) and at the restrictive temperature with diamide (right). (B) Metagene plot (top) and heatmaps (bottom) arranged by decreasing RNA Pol II occupancy of biological replicates of RNA Pol II ChIP-seq for vector, HA-yTBP, HA-yTBPc and HA-mNTD-yTBPc from TSS to TES of diamide-induced genes at both non-diamide-treated condition (Ctrl) and diamide-treated condition (Diamide). (C) GO term identified for diamide-induced genes. Statistics calculated using one-way ANOVA: *** p-value $\leq$ 0.001; **** p-value $\leq$ 0.0001. Non-significant is not shown.

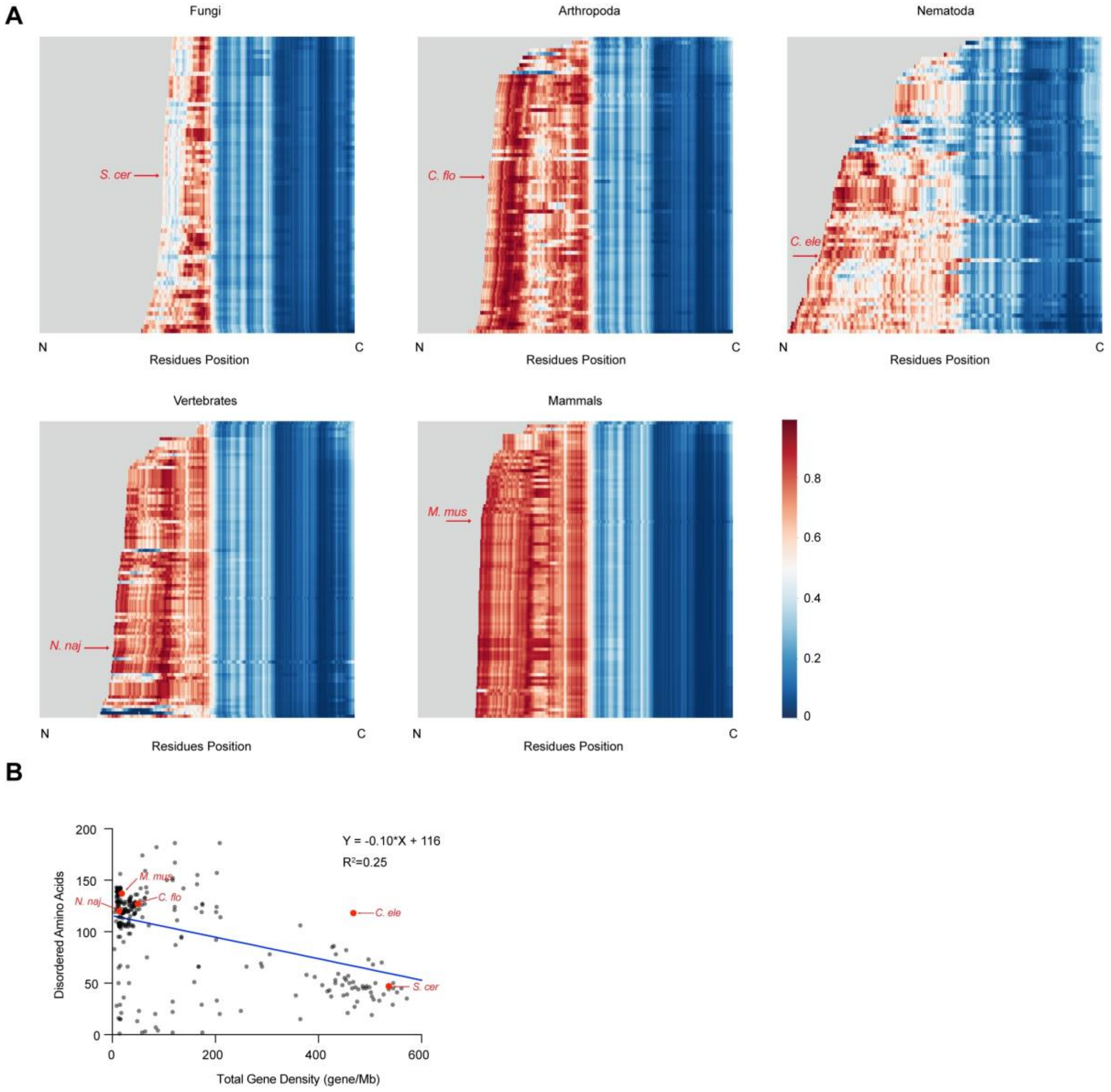


**Figure S6. Individual distribution heatmap for TBP homologs' disordered tendency.**

(A)The distribution heatmap of disorder probability along TBP sequence homologs sorted by length and aligned to C-terminal ends in five taxa (Fungi, Arthropoda, Nematoda, Vertebrates and Mammals). A higher score represents a higher likelihood that the amino acid is in a disordered region. *S. cerevisiae*, *C. floridanus*, *C. elegans*, N. *naja*, and *M. musculus* were labelled as examples from each taxa. (B) The number of predicted disordered amino acids correlates inversely with total gene density.
